# Sorting at ciliary base and ciliary entry of BBSome, IFT-B and IFT-A

**DOI:** 10.1101/2024.03.05.583485

**Authors:** Aniruddha Mitra, Evangelos Gioukakis, Wouter Mul, Erwin J.G. Peterman

## Abstract

Anterograde intraflagellar transport (IFT) trains, composed of IFT-B, IFT-A and BBSome subcomplexes, are responsible for transporting ciliary proteins into the cilium. How IFT subcomplexes reach the ciliary base and assemble into IFT trains is poorly understood. Here, we perform quantitative single-molecule imaging in *C. elegans* chemosensory cilia to uncover how IFT subcomplexes arrive at the base, organize in IFT trains, and enter the cilium. We find that BBSomes reach the base via diffusion where they either associate with assembling IFT trains or with the membrane surrounding the base. In contrast, IFT-B and IFT-A reach the base via directed transport on vesicles that stop at distinct locations near the base. Individual subcomplexes detach from the vesicles into a diffusive pool and associate to assembling trains. Our results indicate that the assembly of IFT trains is a step-wise process involving the subsequent incorporation of first IFT-B, then IFT-A and finally BBSomes.

## Introduction

Primary or sensory cilia are essential, “antenna-like” organelles protruding from most eukaryotic cells to sense the external environment and act as signal transducers^1^. Ciliary structure and function are maintained by a highly regulated transport process, referred to as intraflagellar transport (IFT), where IFT trains, driven by kinesin-2 and IFT-dynein motors, ferry ciliary proteins from base to tip and back again along a microtubule-based axoneme^2,3^. At the base, a ciliary gate, composed of transition fibers and the transition zone (TZ), prevents the diffusive entry and exit of both membrane-bound and soluble proteins^4,5^. To enter, several ciliary proteins have been shown to associate with anterograde IFT trains, assembled at the ciliary base, to be transported through the ciliary gate (Figure 1B). Anterograde IFT trains are ordered polymeric structures (>80MDa) with periodic repeats of IFT-B subcomplexes (each IFT-B complex consisting of 16 proteins) at their core^6^. To this core, IFT-A subcomplexes (9 proteins) attach, positioned away from the axoneme facing the ciliary membrane, with a periodicity suggesting an IFT-B/IFT-A stoichiometry of 2:1^7^. How the third subcomplex, the BBSome (complex of 8 BBS proteins) associates with anterograde IFT trains remains unknown. In recent years, FRAP^8,9^ and structural^10^ studies in *Chlamydomonas*, as well as a single-molecule fluorescence study in *C. elegans* ^11^, have revealed a structural and dynamic picture of how anterograde IFT trains assemble at the base. It appears that IFT trains tether from one end to the TZ while being assembled, with IFT-B forming a scaffold to which IFT-A and IFT dynein attach subsequently from the dendritic side ^10^. Kinesin-2 motors only associate with trains just prior to the trains’ departure, while the cargo tubulin attaches to fully assembled and moving trains^11^. So far, it has not been investigated when and where the BBSome assembles to IFT trains. An additional open question is how IFT components are transported from their sites of synthesis in the soma (via the dendrite in *C. elegans* chemosensory cilia) to the ciliary base and are organized for assembly into trains. In recent years, IFT proteins have been linked to an extraciliary role, in vesicular trafficking^12,13^. Several IFT-B and IFT-A proteins appear to share a common ancestry with classical vesicular coat proteins (COPs)^14,15^, and electron and light-microscopy studies have revealed that some of these proteins associate with periciliary vesicles^16–19^. Because of this, it has been proposed that some IFT components might reach the ciliary base via active microtubule-based transport. A recent FRAP study in *Chlamydomonas*, however, has indicated that disruption of cytoplasmic microtubules does not affect the recruitment of IFT components at the ciliary base^8^. This led the authors to propose that the IFT components form a diffusive pool near the base, with proteins assembling onto trains via a diffusion-to-capture mechanism ^20^. While recent studies provide important insights in the ensemble dynamics of IFT components outside cilia, a clear picture at the single-molecule level is still lacking.

**Figure 1:**
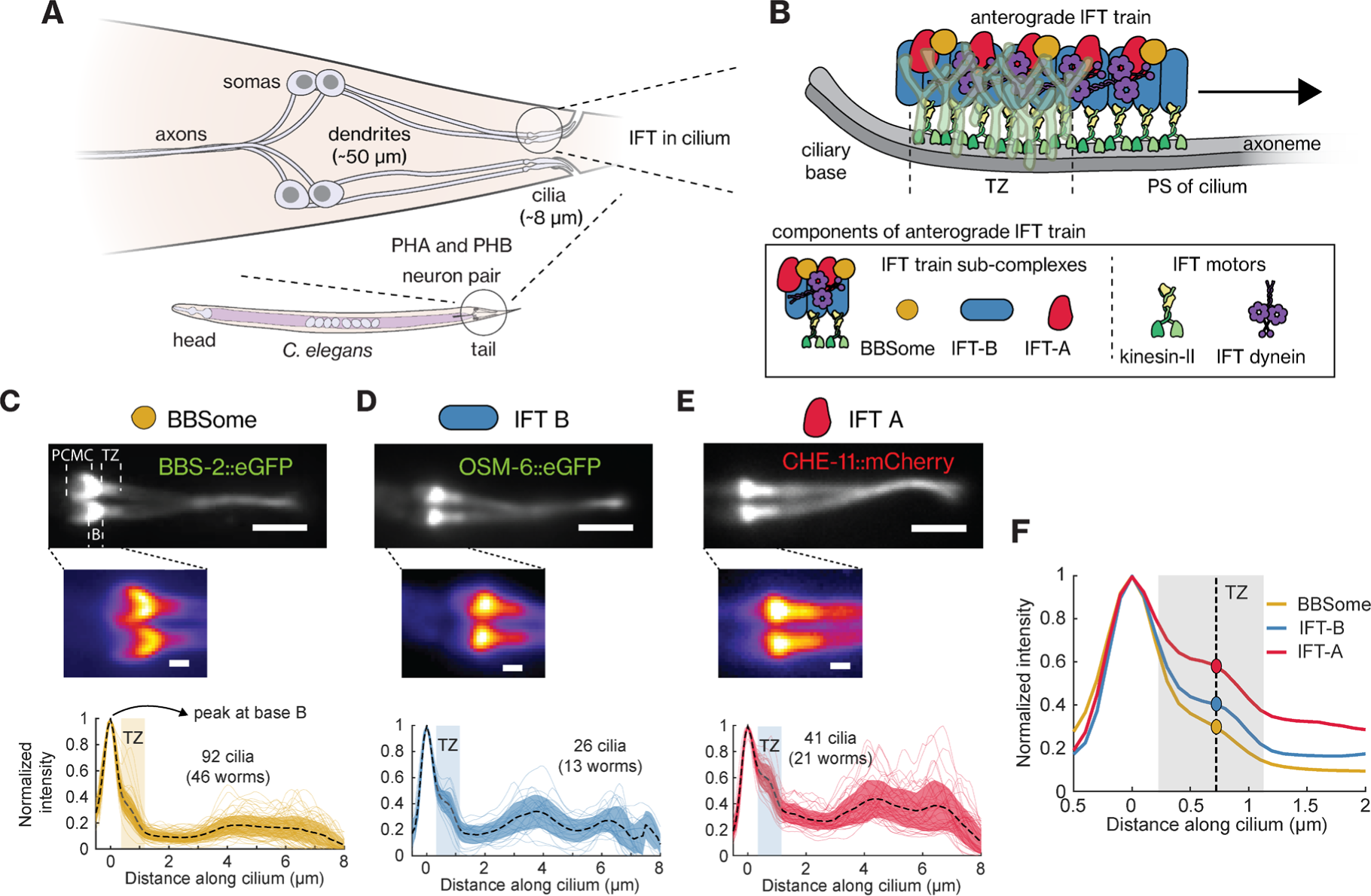
BBSome, IFT-B and IFT-A show distinct localization patterns near the ciliary base of PHA/PHB cilia. **(A)** Schematic diagram of the two pairs of phasmid neurons (PHA/PHB) located at the tail of *C. elegans*. These neurons have ∼50 *μm* long dendrites that connect the somas to the cilia (∼8 *μm* long), with the ciliary tip protruding out of the worm body. **(B)** Top: Illustration of intraflagellar transport (IFT) of an anterograde IFT train crossing the transition zone (TZ; indicated by the Y-shaped linkers (cyan)) to enter the proximal segment (PS) of the cilium. Bottom: Components of an anterograde IFT train. **(C-E)** Representative time**-**averaged fluorescence images (top; corresponding movies in Video S1; insets below show zoom-in of the ciliary base), and the average fluorescence-intensity profiles (bottom; plotting mean ± S.D.) obtained from multiple cilia, for **(C)** BBSome (eGFP::BBS-2, an IFT-train-subcomplex BBSome subunit; 92 cilia from 46 worms), **(D)** IFT-B (OSM-6::eGFP, an IFT-train-subcomplex B subunit; 26 cilia from 13 worms) and **(E)** IFT-A (CHE-11::mCherry, an IFT-train-subcomplex A subunit; 41 cilia from 21 worms). All fluorescence-intensity profiles are normalized to the peak at the ciliary base and the profiles from individual cilia are also plotted in the background. Scale bar, 2 µm. PCMC, periciliary membrane compartment; B, ciliary base; TZ, transition zone. **(F)** The average fluorescence-intensity profile of BBSome, IFT-B and IFT-A, with the shaded area roughly indicating the location of the TZ and the dotted line indicating the centre of the TZ. The relative intensity at the TZ with respect to the peak at the ciliary base is different for the IFT-train-subcomplexes.

In this study, we perform single-particle imaging to visualize the dynamics of individual fluorescently labeled IFT components in the ciliated chemosensory neurons of *C. elegans*. We focus on the PHA/PHB neuron pairs located at the tail of the worms. In these neurons, the cilium is not directly connected to the soma, but via the ∼40-50 *μ*m long dendrite (Figure 1A). Single-particle tracking and analysis reveal how IFT subcomplexes (BBSome, IFT-B and IFT-A) and IFT motors (kinesin-II and IFT-dynein) are transported through the dendrite, from their sites of synthesis in the soma to the periciliary membrane compartment (PCMC; located at the transition between dendrite and cilium). Furthermore, data and analysis reveal a comprehensive picture of the sorting dynamics of individual BBSome, IFT-B and IFT-A subcomplexes in the PCMC as well as their interaction dynamics with anterograde IFT trains assembling at the ciliary base.

## Results

### IFT-train sub-complexes have different localization patterns at the ciliary base

To study IFT-train assembly in the PCMC and at the ciliary base we imaged the ensemble dynamics of BBSome, IFT-A and IFT-B subcomplexes in the chemosensory PHA/PHB neurons of live *C. elegans*. We used BBS-2::eGFP as a marker for the BBSome, OSM-6::eGFP (IFT52 in other organisms) as a marker for IFT-B and and CHE-11::eGFP for IFT-A (IFT140 in other organisms; see Table S1 for a complete list of the strains used in this study). In the following we will assume that these three proteins are always present in BBSome, IFT-B and IFT-A subcomplexes respectively, a point that will be discussed in detail later. Our ensemble fluorescence image sequences showed similar dynamics as observed before for these proteins (Video S1)^21,22^. From these sequences, we obtained time-averaged fluorescence images and intensity profiles showing the ciliary distributions of BBSome (Figure 1C), IFT-B (Figure 1D) and IFT-A (Figure 1E) subcomplexes. The intensity distributions of these subcomplexes are roughly similar, all peaking at the ciliary base, with a shoulder in the TZ, decreasing in the proximal segment, and increasing again in the distal segment, where the two cilia of a PHA/PHB pair overlap. On the dendritic side of the TZ, however, IFT-A and IFT-B appeared to be more restricted to locations close to the ciliary base, while BBSome locations appeared to be extended further towards the PCMC (inset Figures 1C-1E). Furthermore, BBSome and IFT-B subcomplexes appeared to be more specifically enriched at the ciliary base than IFT-A (Figure 1F), which shows a substantially larger shoulder in the TZ. These subtle differences in distributions of the BBSome, IFT-B and IFT-A subcomplexes near the ciliary base suggest that these complexes are sorted differently before entry into the cilium.

### BBSomes reach the ciliary base by diffusion, while IFT-B and IFT-A are actively transported along the dendrite

In the PHA/PHB sensory neurons of young adult worms, the cilia are separated from the soma by ∼40 − 50 *μm* long dendrites (Figure 1A). To investigate how individual IFT components synthesized in the soma are transported along the dendrite to reach the ciliary base, we visualized the dynamics of individual molecules in the PHA/PHB dendrites using small-window illumination microscopy (SWIM)^23^. The idea of SWIM is to excite and photobleach only a small region of the sample (5-15 *μm* diameter), allowing continuous and long-term entry of “fresh” proteins, which have not yet been photobleached, into the excited region. Using this approach, we observed that BBSome subcomplexes diffuse rapidly in the dendrite (Figure 2A and Video S2), similar to what we have observed before for IFT dynein and kinesin-II (Figure S1A)^11^. From these image sequences, we extracted single-particle trajectories, which were used to calculate the diffusion coefficients using a covariance-based estimator^24^. We found that BBSome subcomplexes diffuse faster than IFT dynein but slower than kinesin-II (Figures 2B-2C and Figures 1B-1C; overview of the statistics in Table S2). This was expected given that the diffusion coefficient scales with the inverse of the cube root of the mass^25^, and the BBSome, IFT-dynein and kinesin-II have masses of ∼0.5 MDa, ∼1.5 MDa and ∼0.3 MDa, respectively. Furthermore, this supports our assumption that BBS-2 is incorporated in the BBSome subcomplex and is not diffusing on its own, which would result in a much larger diffusion coefficient given its mass of only ∼80 kDa. Our observations suggest that the BBSome and IFT motors do not use active transport for their journey from soma to cilium along the dendrite but utilize free diffusion. It can be estimated (< *t* >≈ *δx*^2^/2*D*) that transport along the whole length of the ∼50 *μm* long dendrite would then take on average ∼13 mins (kinesin-II) to 42 mins (IFT-dynein).

**Figure 2:**
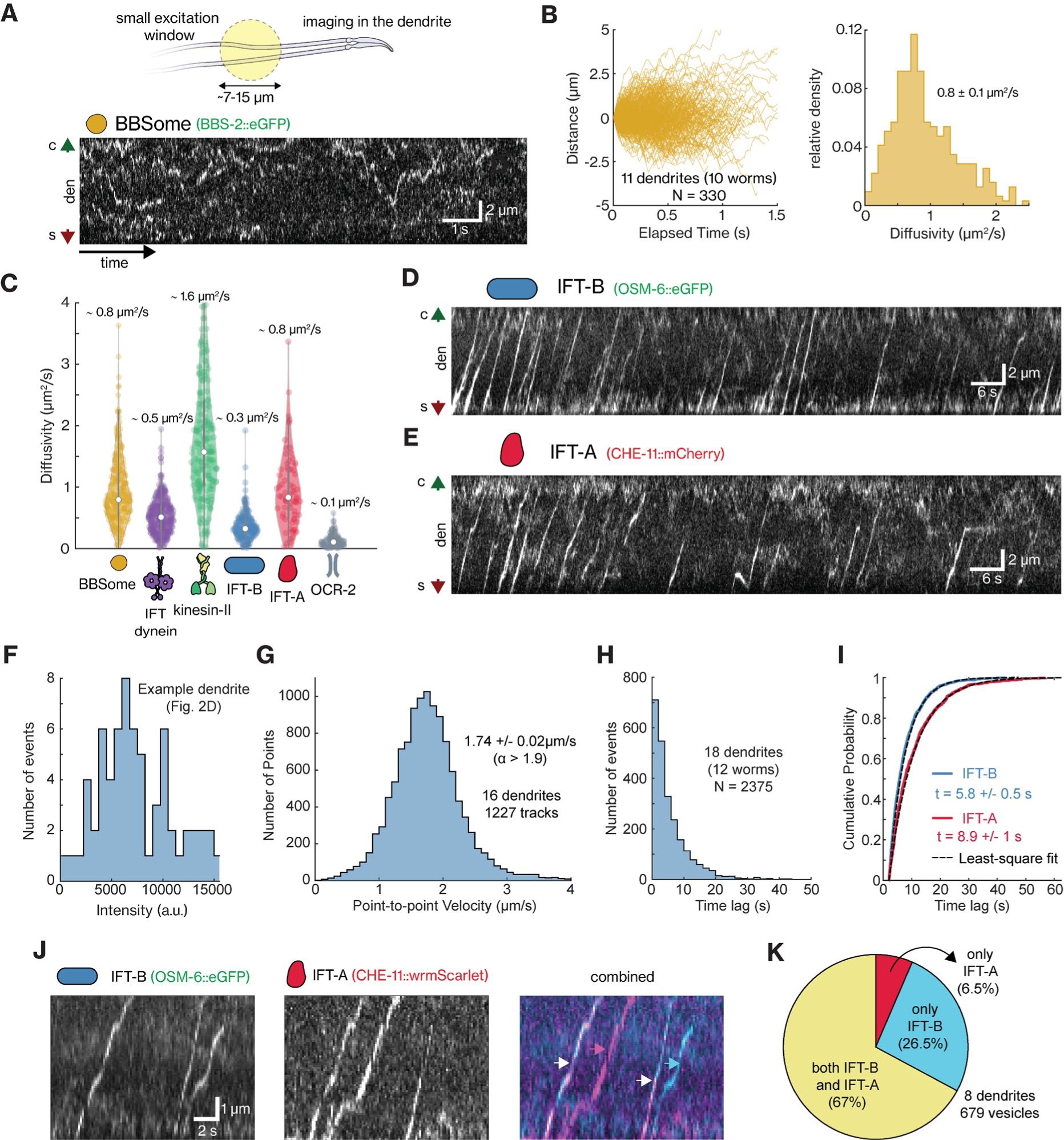
In the dendrite, BBSome and IFT motors are transported diffusively while IFT-A and IFT-B are transported in a directed manner. **(A)** Top, illustration of small-window illumination microscopy (SWIM) that allows single-molecule imaging of IFT components in the dendrites of PHA/PHB neurons. Bottom, representative kymograph (space-time intensity plot) of eGFP::BBS-2 indicates that single BBSome complexes diffuse across the dendrite (see Video S2). Green and red arrows indicate the cilium (c) and soma (s) directions, respectively. **(B)** Left, distance-time plots of all tracked BBSome particles diffusing in the dendrite (157 tracks from 9 dendrites). Right, histogram of the diffusivity of individual tracks, estimated using a covariance-based estimator (CVE), with the average diffusivity 0.8 ± 0.1 *μm*^2^/*s*. **(C)** Violin plot of the diffusivity of BBSome, IFT-B, IFT-A, IFT dynein, kinesin-II and OCR-2-associated vesicles in the dendrite. Each point represents the diffusivity of a tracked diffusive particle, estimated using CVE. The median and interquartile range is indicated in the violin plots. **(D-E)** Representative kymographs of IFT-B (OSM-6::eGFP; D) and IFT-A (CHE-11::mCherry; E) in dendrites display that these complexes are transported in a directed manner in packets from the soma towards the cilia (see Video S3). Green and red arrows indicate the cilium (c) and soma (s) directions, respectively. **(F)** Histogram of intensity of directed packets of IFT-B imaged in example dendrite shown in 2D and Video S3 (left). **(G)** Histogram of point-to-point velocities obtained from the parts of the 1227 tracks (16 dendrites) of OSM-6 packets that were highly directed (*⍺* > 1.9), with the average velocity 1.74 ± 0.02 *μm*/*s*. **(H)** Histogram of the time lag between subsequent OSM-6 packets in 18 dendrites (N = 2375). **(I)** The time-lag distribution of OSM-6 (blue) and CHE-11 (red) packets plotted as a cumulative distribution function, overlayed with the least square fit to the function. The characteristic time between subsequent OSM-6 packets is 5.8 ± 0.5 s and between subsequent CHE-11 packets is 8.9 ± 1 s. Average value and error are estimated using bootstrapping (see Methods). **(J)** Representative kymograph obtained from dual-colour imaging of IFT-B (OSM-6::eGFP) and IFT-A (CHE-11::wrmScarlet) in dendrites. Left: IFT-B, middle: IFT-A, right: combined. Right: Colour of the arrows indicate the vesicle composition (cyan: only IFT-B, magenta: only IFT-A and white: both). **(K)** Pie-chart displaying that 6.5% of vesicles contain only IFT-A, 26.5% contain only IFT-B and 67% contain both IFT-B and IFT-A (679 tracks from 8 dendrites).

We next investigated how IFT-B and IFT-A subcomplexes are transported along the dendrite towards the ciliary base. We observed that both IFT-B and IFT-A, in contrast to the other IFT components, are actively transported in a directed manner from soma to ciliary base (Figures 2D-2E, Video S3). Judging from the intensities of the fluorescence spots, multiple IFT-B or IFT-A subcomplexes are transported together, possibly as IFT-B and IFT-A -coated vesicles, since several IFT-A and IFT-B proteins have been shown to be structurally similar to COP-I, II and clathrin proteins^14,15^ and take part in vesicle trafficking outside cilia^12^. Apart from these vesicles, we also observed that a fraction of the subcomplexes moved diffusively. The diffusion coefficients for this diffusing IFT-B (∼2.2 MDa) and IFT-A (∼0.8 MDa) subcomplexes are in line with the trend estimated from their molecular weights (Figure 2C and Figure S1B-S1C; statistics in Table S2). The IFT-B diffusion coefficient is smaller than that of the other IFT components, but significantly higher than that of OCR-2-containing dendritic vesicles (ciliary TRPV-channel protein; Figure S1B-S1C; also reported previously^23^). This could indicate that individual IFT-B and IFT-A subcomplexes diffuse, while vesicles containing IFT-B and IFT-A subcomplexes are actively transported. We observed that the fluorescence intensity of these vesicles differed considerably (Figure 2F and Figure S2A), indicating that their size and/or protein content is variable. To obtain quantitative insight in the vesicle transport dynamics, we extracted individual trajectories. Tracked events showed either mostly directed motions, or bouts of directed motion, interspersed with diffusive episodes and pauses (Figure 2D-2E). To extract the mode of transport, we subjected the trajectories to a windowed mean squared displacement (MSD) approach ^11,26,27^, which allows determination of the anomalous exponent (α). For directed motion, α is expected to be equal to 2; for diffusive motion, α = 1; and for sub-diffusion or pausing α < 1. For IFT-A and IFT-B trajectories that appeared completely directed, we indeed found α values of ∼2, while for other trajectories displaying more varied motility, we found α values between 1.5 and 2. After filtering the complete data sets for directed motion (only taking into account time windows when α > 1.9), we determined that the average point-to-point velocity of directed transport is 1.74 ± 0.02 μm/s for IFT-B (average ± error estimated using bootstrapping; Figure 2G and Figure S2B) and 1.94 ± 0.02 μm/ for IFT-A (Figure S2C). In the dendrites of *C. elegans* PHA and PHB neurons –where the microtubules are oriented with their minus-ends pointing in the direction away for the soma, towards the cilium^28^– all anterograde dendritic transport is expected to be driven by the minus-end motor cytoplasmic dynein-1. The differences we observed in the velocities of IFT-B and IFT-A-coated vesicles are consequently most likely not caused by differences in the nature of the motor driving transport, but by the number of motors engaged, or other properties of the vesicles, like size and membrane properties, or variations in the local architecture of microtubules travelled by the different vesicles. We also considered the time intervals between subsequent vesicles and found that they are exponentially distributed (Figure 2H and Figure S2D) with a characteristic time of 5.8 ± 0.5 s for IFT-B and 8.9 ± 1 s for IFT-A (Figure 2I). Before, we have shown that the ciliary transmembrane protein OCR-2, also associates with vesicles showing similar transport dynamics in the dendrite. A key difference with IFT-particle subcomplex vesicles is that the characteristic time between subsequent OCR-2 vesicles is substantially longer (19.8 ± 3.2 s)^23^. Finally, we imaged IFT-B and IFT-A simultaneously (Figure 2J, Video S4; OSM-6::eGFP and CHE-11::wrmScarlet) and observed that ∼67% of the vesicles contain both IFT-B and IFT-A, ∼26.5% only IFT-B and ∼6.5% only IFT-A (Figure 2K). This suggests that IFT-B and IFT-A subcomplexes are mostly co-transported by the same vesicle but that a substantial minority of vesicles is loaded with only IFT-B, which can explain the difference in time intervals between IFT-B and IFT-A-coated vesicles observed in Figure 2I. Taken together, these findings indicate that IFT-A and IFT-B subcomplexes, and transmembrane proteins are actively transported, attached to vesicles moving along the dendrite towards the cilium. In contrast, other protein complexes, including the BBSome and the IFT motors, move diffusively along the dendrite.

### BBSomes bind to the PCMC membrane or to assembling anterograde IFT trains

We next used SWIM to investigate the dynamics of individual IFT-train subcomplexes in the region where the ciliary base meets the dendrite (Figure 3A left), focusing first on BBSomes (Figure 3A right and Video S5). The BBSome tracks showed three distinct kinds of behaviors. (I) In some tracks, BBSomes switched from a diffusive to an immobile state close to the ciliary base, followed by a short pause, before they started to move into the cilium, speeding up beyond the TZ (Figure 3C left). These tracks are similar to those we observed before for IFT motors entering the cilium^11^ and we infer that they represent BBSome subcomplexes that bind to immobile, assembling IFT trains at the ciliary base^10^ and are transported into the cilium after train assembly has completed. We refer to these tracks as “ciliary-entry events”. (II) In other tracks, BBSomes switched from a diffusive to an immobile state apparently binding to the PCMC membrane, in most cases close to the ciliary base (Figure 3C middle). In these tracks, we did not observe ciliary entry, but we often saw BBSomes being released again from their static location and diffusing in the dendrite. In other cases, BBSomes appeared to start diffusing again while remaining attached to the PCMC membrane. These latter particles appeared to diffuse substantially faster and often “hopped” to another static location in the PCMC. We refer to this second class of track as “PCMC-binding events”. It has been shown before that BBSomes can be recruited from an autoinhibited state in solution to a membrane-bound state by ARF-like GTPase ARL6/BBS3 ^29–31^. We thus propose that the PCMC-binding events represent BBSomes freely diffusing in the dendrite that switch conformation and bind to the PCMC membrane, likely interacting with ARL6, which has been shown to be enriched in the membrane surrounding the ciliary base in mammalian cells^32,33^. (III) In the few retrograde tracks we observed, BBSomes appeared to exit the cilium and subsequently remained stuck in the PCMC (Figure 3C right).

**Figure 3:**
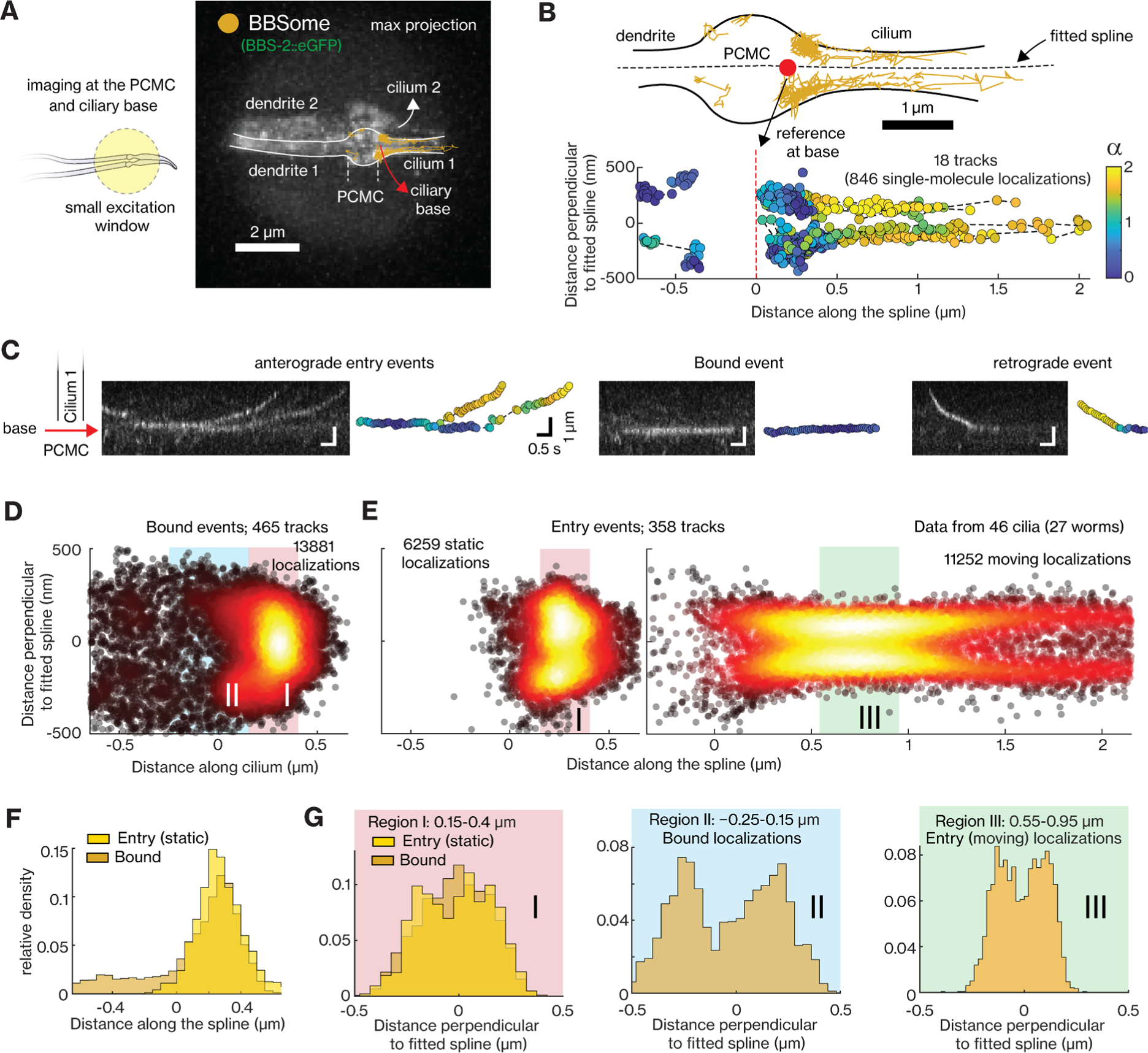
Analysis of the dynamics of individual BBSomes at PCMC and proximal part of cilium. **(A)** Right, maximum projection of an example movie (Video S5) shows the section of dendrites, PCMC and cilia of a PHA/PHB neuron pair illuminated by a small excitation window (left, illustration), where the dynamics of eGFP::BBS-2 (BBSome) is visualized. **(B)** Top, tracked single-molecule events with the outline of the cell indicated. Dotted black line displays the fitted spline and the red point indicates the reference point at the base. Bottom, the tracked events plotted in ciliary coordinate. The colour of the localizations indicate the *⍺*-value, a measure of degree of directedness of the event at that localization. *⍺*-value is ∼2 for purely directed transport and ∼1 for purely diffusive transport. **(C)** Kymographs displaying example anterograde entry events (left), event bound at the ciliary base (middle) and retrograde event (right) with corresponding tracks (colour of localizations indicate the *⍺*-value). **(D-E)** Super-resolution map of single-molecule localizations obtained 465 tracks of PCMC-binding events (D; N = 13881 localizations) and 358 tracks of entry events (E; left panel: N = 6259 static localizations (*⍺* < 1); right panel: N = 11252 moving localizations (*⍺* > 1.2)) obtained from 46 cilia. **(F)** Distribution of distance along the cilium for localizations of PCMC-binding events and static localizations of entry events. **(G)** Distribution of distance perpendicular to ciliary spline for localizations of PCMC-binding events and static localizations of entry events (*⍺* < 1) between 0.15-0.4 *μm* (left; region I), bound events between −0.25-0.15 *μm* (center; region II) and moving localizations of entry events between 0.55-0.95 *μm* (right; region III). Regions I, II and III are indicated in D-E.

To obtain a more detailed spatial picture of static BBSome localizations at the ciliary base, we extracted super-resolution fluorescence maps from the tracks of the PCMC-binding and ciliary entry events (Figures 3D and 3E left, respectively; statistics in Table S3). To this end, we filtered the tracks using the windowed MSD-approach for α <= 1, (see example tracks in Figure 3B). The distribution of static localizations from the PCMC-binding tracks shows a maximum at the ciliary base (at ∼0.3 *μm*), but with a long tail into the dendrite (Figure 3E). The distribution of static localizations from the ciliary-entry events also showed a sharp peak, slightly shifted towards the dendrite (at ∼0.2 *μm*; Figure 3E) but without the long tail into the dendrite. The distribution of the moving localizations of BBSome entry events (α > 1.2; Figure 3E right) looked similar to those we observed before for the IFT motors^11,34^, revealing the characteristic shape of the initial part of the cilium: relatively broad at the ciliary base, tapering at the TZ and “bulging” out at the proximal segment of the cilium. We next looked into the distributions perpendicular to the ciliary long axis. For localizations of ciliary-entry as well as PCMC-binding events in region II (Figure 3G), axial distributions show two maxima equidistant from the centre. Before^11^, we have interpreted the bimodal distributions to be due to the 2D projection (by our imaging approach) of the 3D distribution along a hollow cylinder (the axoneme or the PCMC membrane), see also Figure S3B. For the PCMC-binding events in region I (close to the ciliary base) it is likely that the narrow membrane width in this region does not allow us to discern the two maxima and only a single maximum at the centre is observed (Figure 3G left). Taken together, single-particle tracking reveals that BBSome subcomplexes show two distinct behaviours at the ciliary base after exiting the dendrite: one population of BBSomes docks to the PCMC membrane, primarily close to the ciliary base, while the other attaches to assembling IFT trains at the ciliary base. The overall single-particle localization maps are consistent with the ensemble intensity profiles at the PCMC and proximal part of the cilia (Figure 1F).

### Directed vesicles of IFT-A and IFT-B are sorted differently at the PCMC and the ciliary base

Next, we imaged the dynamics of IFT-B arriving from the dendrite into the PCMC (see example neuron-pair in Video S6A and tracked single-molecule localizations from one neuron in Figure 4A). We observed that IFT-B-coated vesicles moved directedly and apparently actively from dendrite towards the PCMC, where they slowed down or paused briefly between −1.1 *μm* to −0.7 *μm* from the ciliary base (example events in Video S6B and Figure 4B). Part of the vesicles appeared to continue their directed route, docking at the ciliary base, where they paused, often for several seconds, occasionally hopping from one location to another. Some vesicles did not seem to reach the ciliary base, because the fluorescence signal disappeared at the pause location close to the dendrite-PCMC junction, either by photobleaching or by IFT-B subcomplexes dissociating from the paused IFT-B-coated vesicles and diffusing away (as could be observed occasionally: Figure 4C and Video S6B). Collectively, we refer to these events as “vesicle” events. We also observed diffusive IFT-B docking at the ciliary base, pausing for a while, before entering the cilium (Figure 4C). This behavior is characteristic of “entry events”: IFT components associating with an assembling IFT train that enter the cilium after the train starts moving (like we have observed before for BBSomes (Figure 3C) and other IFT components^11^). Remarkably, we did not observe direct entry of IFT-B subcomplexes from coated vesicles that had been transported towards the ciliary base and paused there. It might be that such entry events are rare, since most complexes photobleach during the relatively long pause at the base, or since other vesicles or diffusive complexes arrive in the vicinity, making it difficult to distinguish them reliably as single events. Apart from vesicle and entry events, we also obtained “stuck” events, which are due to diffusive particles docking at the ciliary base, where they remain static until the signal disappears due to photobleaching or gradual dissociation of IFT-B.

**Figure 4:**
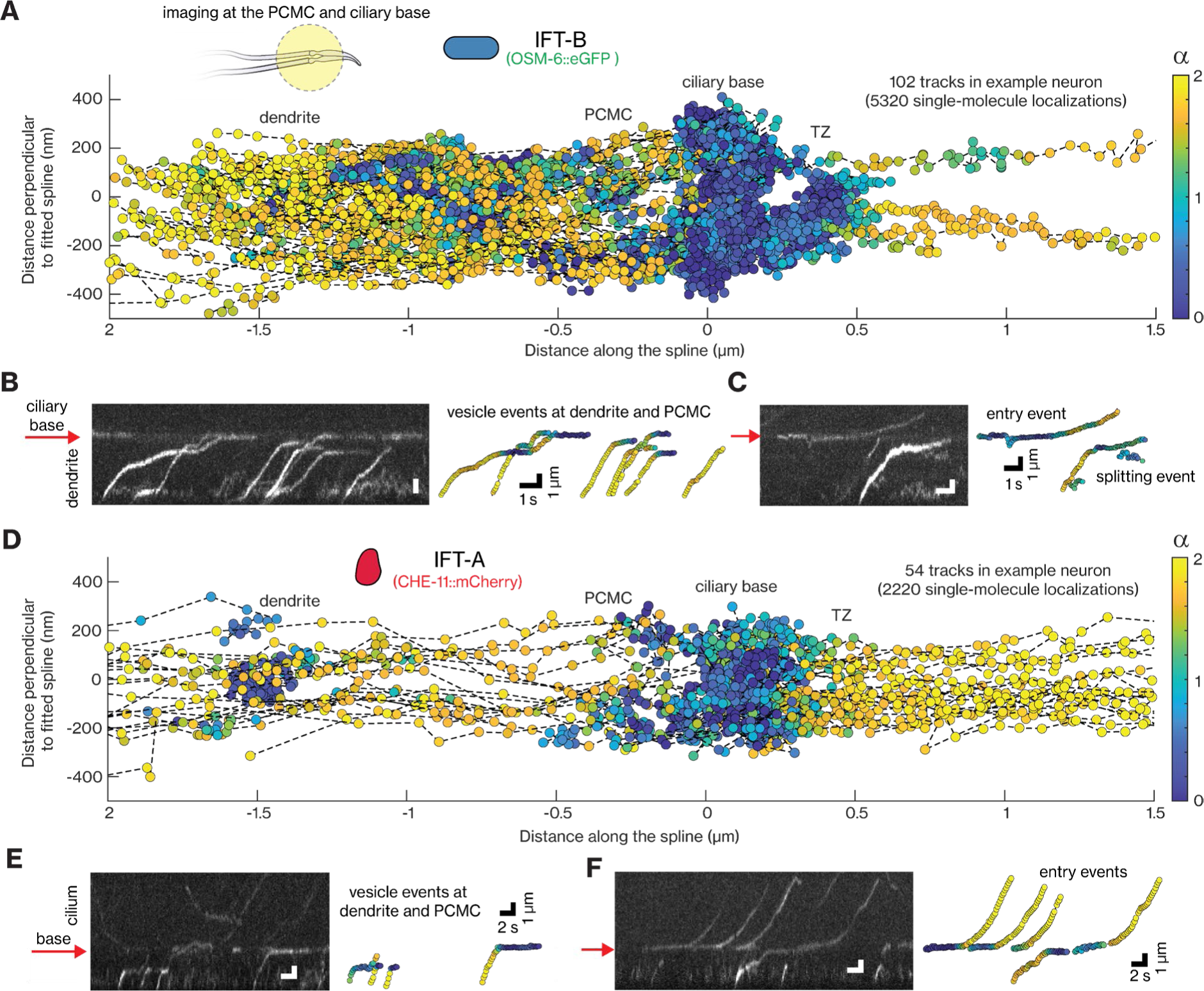
Dynamics of individual IFT-B and IFT-A complexes at PCMC and ciliary base. **(A)** Single-particle IFT-B tracks obtained from an example OSM-6::eGFP labelled worm imaged using SWIM (Video S6) plotted in cilia-coordinates. Inset: Illustration of the section of dendrites, PCMC and cilia of a PHA/PHB neuron pair illuminated by a small excitation window (left, illustration), where the dynamics of IFT-B and IFT-A are visualized. **(B-C)** Representative kymograph showing example IFT-B events (left) and corresponding tracks (right). **(B)** Directed IFT-B-coated vesicles moving from the dendrite into the PCMC. These vesicles slow down or pause at the dendrite-end of the PCMC (∼1 *μm* from ciliary base), with individual IFT-B complexes eventually bleaching, diffusing, or reaching the ciliary base in a directed manner where they pause for a much longer time. **(C)** Event on left: An IFT-B complex, diffusing in the dendrite, docks at the ciliary base, hops to another location at the base and enters the cilium after a pause. Event on right: A high-intensity IFT-B-coated vesicle moving from the dendrite towards the cilia, releasing IFT-B complexes that diffuse. **(D)** Single-particle IFT-A tracks obtained from an example CHE-11::mCherry labelled worm (Video S7) plotted in cilia-coordinates. **(E-F)** Representative kymograph showing example IFT-A events (left) and corresponding tracks (right). **(E)** Directed IFT-A-coated vesicles moving from the dendrite into the PCMC. While most IFT-A complexes bleach or switch to a diffusive state as the IFT-A-coated vesicles move through the PCMC, some reach the ciliary base where they pause for a long duration (rightmost event). **(F)** IFT-A complexes, diffusive in the PCMC, dock at the ciliary base and enter the cilium. Occasionally IFT-A complexes reaching the base coated on directed vesicles directly enter the cilia (rightmost event). Red arrow in all kymographs indicate the location of the ciliary base. Scale bar of the kymograph (left) are the same as the scale bar for the corresponding tracks (right). Colour of localizations in all plots indicate the *⍺*-value.

Also, IFT-A displayed rich dynamics near the PCMC and the ciliary base (Figure 4D and Video S7A), with some key differences to IFT-B. IFT-A-coated vesicles did not seem to pause near the dendrite-PCMC junction but slightly before, towards the dendrite, mostly around −1.5 *μm* from the base. Furthermore, a large fraction of IFT-A did not seem to reach the ciliary base. While this is partly due to photobleaching, it appeared that IFT-A dissociates from paused as well as moving vesicles more readily than IFT-B (Figure 4E). IFT-A entry events, where IFT-A subcomplexes from a cytosolic pool dock at the ciliary base and, after a short pause, move into the cilia, were more frequently observed than in the case of IFT-B (Figures 4A, 4D-4E). Partly, this may be due to vesicles containing much fewer IFT-A by the time they reach the ciliary base, making the ciliary base far less busy and single particles far more straightforward to discern. Furthermore, on rare occasions we also observed ciliary-entry events arising directly from an IFT-A-coated vesicle parked at the ciliary base (Figure 4F), suggesting that it is possible for IFT-A to swiftly associate with assembling IFT trains from coated vesicles.

To obtain a more detailed spatial picture of IFT-A and IFT-B arriving at the ciliary base, we generated super-resolution fluorescence maps from the localizations in IFT-B (Figure 5A) and IFT-A (Figure 5B) in vesicle and ciliary entry tracks obtained from multiple cilia and worms, using our windowed MSD approach to filter for static (α < 1) and moving (α > 1.2) localizations (statistics in Table S3). The distribution of static localizations of entry events peaks closer to the TZ (peak at ∼0.2 *μm*) than that of static localizations of vesicle events (peak at ∼0 *μm*), both for IFT-B (Figure 5C) and IFT-A (Figure 5D). This indicates that vesicles stop moving at slightly different locations close to the ciliary base where single IFT-B and IFT-A subcomplexes associate with assembling IFT trains. The distribution of localizations of stuck events is in between those of static entry and vesicles localizations (distributions of IFT-B events in Figure S4A), suggesting that they are a combination of entry events that photobleach before the train has moved into the cilium and diffusive IFT-B-coated vesicles binding at the ciliary base. We observed a second peak in the distribution of static vesicle localizations at the dendrite-PCMC junction for IFT-B at ∼-0.9 *μm* and slightly further in the dendrite for IFT-A at ∼-1.5 *μm* (Figure 5E), as is also evident in the example trajectories shown in Figure 4. Also, the distributions of moving IFT-B and IFT-A vesicle localizations show secondary peaks at ∼-0.9 *μm* and ∼-1.5 *μm*, respectively (bottom panel Figures 5A-5B). Taken together, these super-resolution maps indicate that IFT-A and IFT-B-coated vesicles heading toward the ciliary base initially pause briefly at distinct locations, with IFT-B vesicles primarily pausing near the dendrite-PCMC junction (at ∼-0.9 *μm*) and IFT-A vesicles pausing throughout the dendritic region near the PCMC and at the PCMC (peaking at ∼-1.5 *μm*). Further support for this can be found in the local, averaged point-to-point, axial velocities (Figure 5F), which show that IFT-A-coated vesicles slow down earlier (from ∼-2 *μm*) than IFT-B-coated vesicles (from ∼-1.5 *μm*). Furthermore, the axial distributions of moving IFT-A and IFT-B in the PCMC, appear hollow with substantially more localizations along the periphery than in the center (bottom panel Figures 5A-5B), indicating that the microtubules extending from the dendrite into the PCMC, along which the IFT-coated vesicles are transported, are organized along the PCMC membrane. Overall, our observations reveal that IFT-B- and IFT-A-coated vesicles are differently sorted, pausing at different locations before and at the ciliary base. At these pause locations, IFT-B and IFT-A subcomplexes gradually dissociate from the vesicles, forming a diffusive pool in the PCMC. It is mostly IFT-B and IFT-A subcomplexes from this diffusive pool that associate with assembling IFT trains at the ciliary base. Once assembly of these trains is completed, they ferry IFT-B and IFT-A subcomplexes into the cilium.

**Figure 5:**
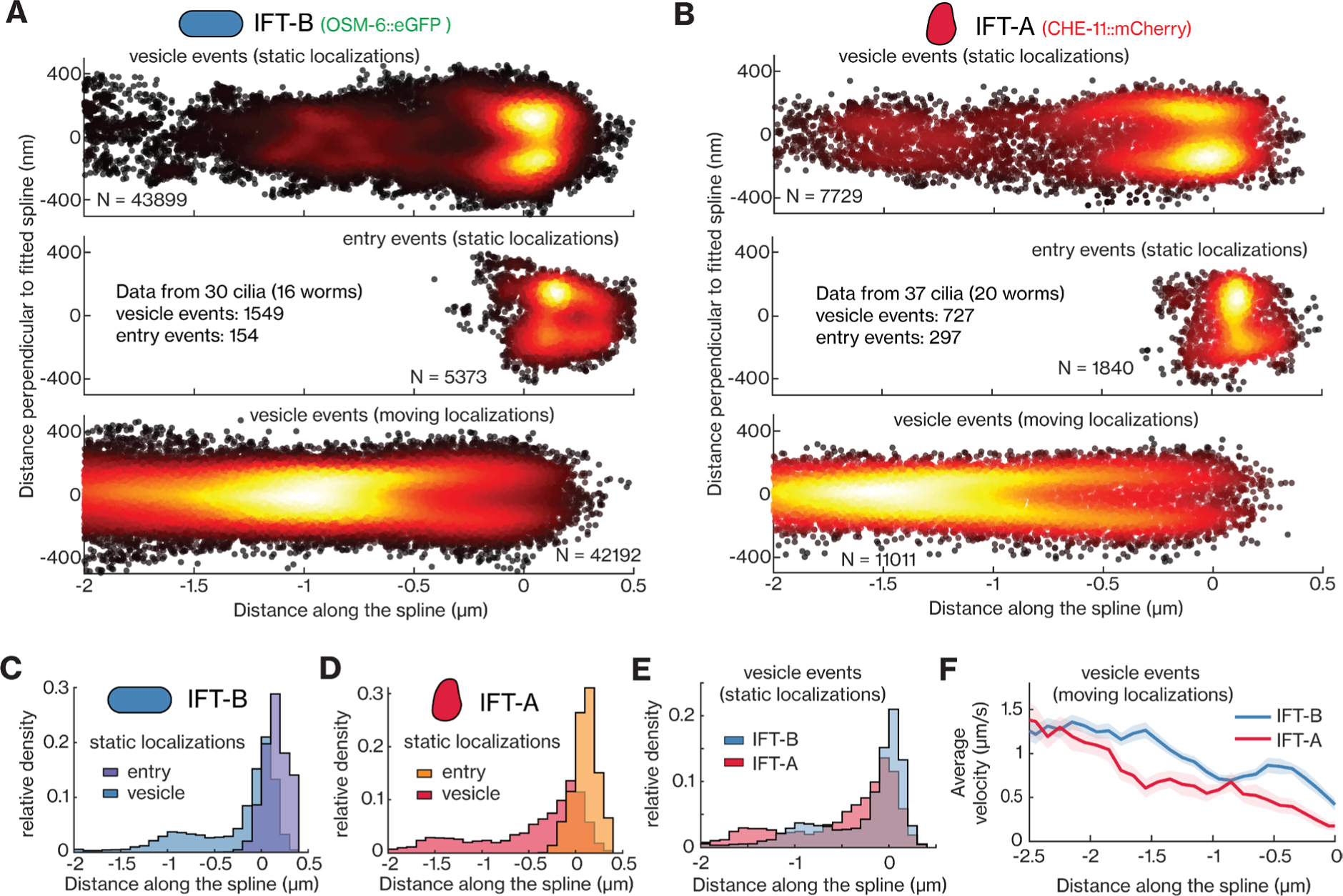
Single-particle localizations reveal that IFT-B and IFT-A are sorted differently in the PCMC and ciliary base. **(A-B)** Super-resolution map of single-particle localizations obtained from tracks of IFT-B and IFT-A. **(A)** Localizations of IFT-B from 30 cilia. Top: static localizations from vesicle events (43899 localizations from 1549 tracks); middle: static localizations from entry events (5373 localizations from 154 tracks); bottom: moving localizations from vesicle events (42192 localizations). **(B)** Localizations of IFT-A from 37 cilia. Top: static localizations from vesicle events (7729 localizations from 727 tracks); middle: static localizations from entry events (1840 localizations from 297 tracks); bottom: moving localizations from vesicle events (11011 localizations). **(C-E)** Distributions of distance along fitted spline for static localizations: **(C)** from vesicle and entry events of IFT-B; **(D)** from vesicle and entry events of IFT-A; **(E)** from vesicle events of IFT-B and IFT-A. **(F)** Velocity distribution between −2.5-0 *μm* along the fitted spline, obtained from moving localizations of vesicle events, for IFT-B (blue) and IFT-A (red).

### BBSome, IFT-B and IFT-A show different dynamics before entering the cilium

Finally, we compared BBSome, IFT-B and IFT-A entry events (left panel Figures 6A-6C). In all these tracks we observed the IFT components, mostly part of a diffusive pool in the PCMC, dock at the ciliary base, presumably by attaching to assembling IFT trains, to pause for a brief interval of time, before moving into the cilium (left panel Figures 6A-6C). We found that the distributions of docking locations are different for the different IFT-train sub-complexes: while most BBSomes dock onto IFT trains between 0-0.4 *μm* from the ciliary base (average 0.19 ± 0.03 *μm*), some events start deeper inside the TZ (0.5 - 1 *μm*). In contrast, IFT-A and IFT-B dock almost exclusively between 0 and 0.3 *μm* (averages 0.10 ± 0.03 *μm* and 0.07 ± 0.03 *μm*, respectively (middle panel Figures 6A-6C)). Similar to this observation, the distributions of static localizations of BBSome, IFT-B and IFT-A entry events also highlight that, on average, BBSomes are statically associated with IFT trains slightly closer to the TZ than IFT-B and IFT-A (Figure S4). For each entry track we also determined the measured pause time, *t*_*p*_*m*_, defined as the duration it takes a single complex to move 100 nm along the cilium. We found that *t*_*p*_*m*_ is significantly smaller for BBSomes (0.8 ± 0.1 s, average ± error estimated using bootstrapping; right Figure 6A) than for IFT-A (1.2 ± 0.2 s; right Figure 6C), and IFT-B (1.8 ± 0.4 s; right Figure 6B). We note that these measured pause durations are highly affected by eGFP/mCherry photobleaching, which terminates many trajectories before they can be classified as entry events. To account for this we use simulations (see Figure S5A and methods for details; as performed before for estimating pause times of IFT motors^11^). These simulations allow us to estimate the actual pause times of BBSomes to be ∼0.4-0.7 s, of IFT-B >9 s (Figure S5F; example simulation in Figure S5E), and of IFT-A ∼2-11 s (Figure S5H; example simulation in Figure S5G; we note here that the estimation of the actual pause duration becomes relatively inaccurate when the measured pause time is similar to the characteristic photobleaching time). These estimates reveal that BBSomes pause for, on average, a much shorter duration than IFT-A and IFT-B subcomplexes before ciliary entry. From the moving localizations in these entry tracks, we obtained the average point-to-point velocity profiles, which were very similar for the three IFT train complexes (Figure 6D) as well as IFT dynein^11^, with the velocity increasing to ∼0.4 *μm*/*s* in the first ∼0.6 *μm* of the TZ, then remaining constant until ∼1.1 *μm* before increasing steeply after ∼1.1 *μm*, where the TZ ends. Finally, we estimated the diameter of the 3D cylinder along which the moving IFT components are distributed and found that BBSomes and IFT-A subcomplexes localize further away from the longitudinal axis of the ciliary axoneme than IFT-B (Figure 6E). This observation correlates with studies that reveal that IFT-A and BBSome interact with ciliary membrane proteins^1,35^ and would structurally require to face the ciliary membrane. IFT-B, on the other hand, was located closer to the axoneme, to which it is connected via kinesin-2 motors. Thus, our observations indicate that IFT-B and IFT-A associate with assembling IFT trains during the early stages of assembly when they are immobile, while BBSomes associate with IFT trains at the final stages of assembly or already assembled trains moving inside the cilium. Furthermore, IFT-train components have the same characteristic velocity profile since they move together across the TZ to enter the cilium. Single-molecule localization maps hint towards close association of the BBSome and IFT-A with the ciliary membrane.

**Figure 6:**
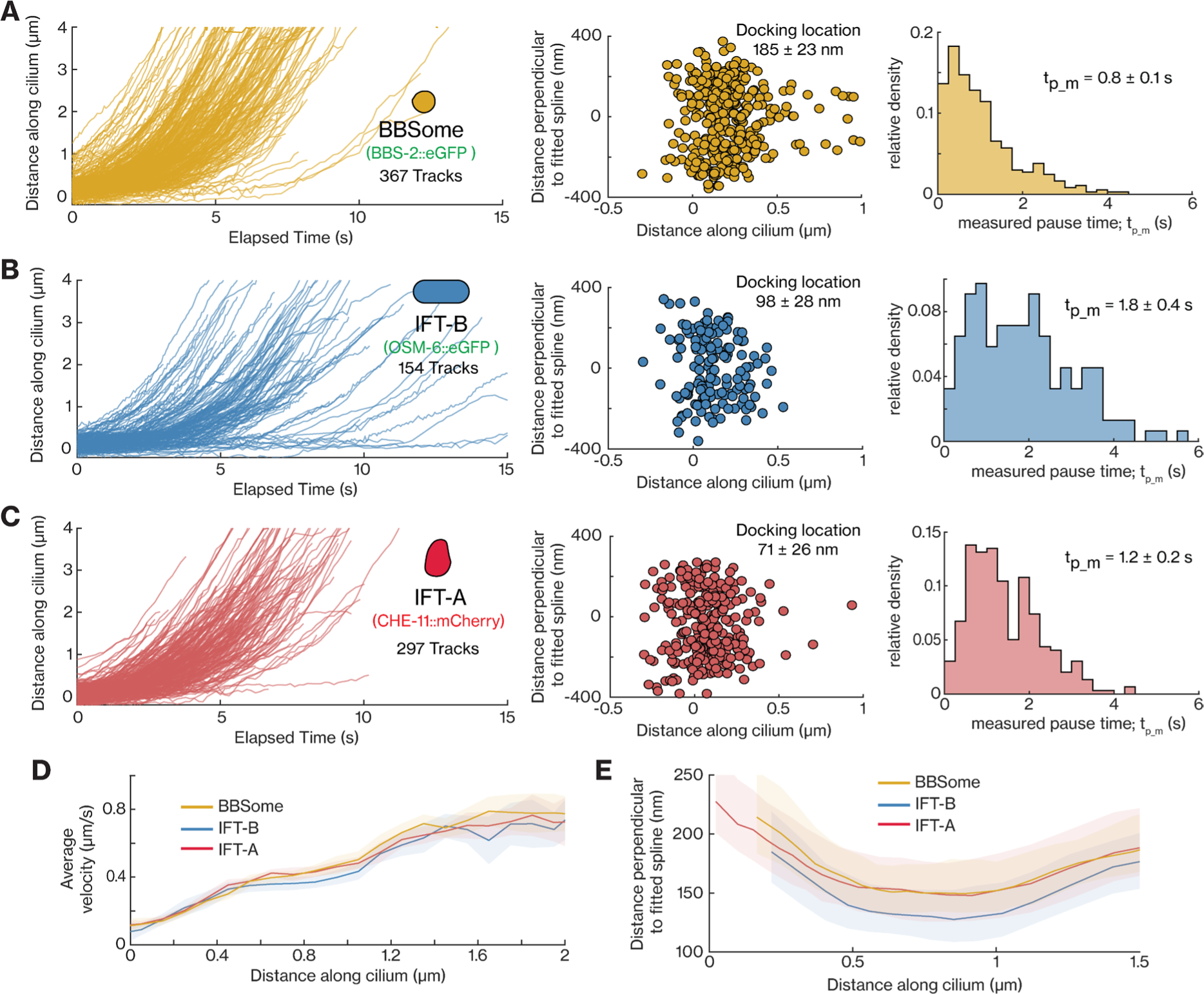
Dynamics of individual BBSome, IFT-B and IFT-B complexes entering cilia. **(A-C)** Distance-time plots (left), distribution of docking locations (middle) and distribution of measured pause time (right) for IFT train complexes: **(A)** 367 entry tracks of BBSome (BBS-2), average docking location 185 ± 23 nm, average measured pause time is 0.8 ± 0.1 s; **(B)** 154 entry tracks of IFT-B (OSM-6), average docking location 98 ± 28 nm, average measured pause time is 1.8 ± 0.4 s; **(C)** 297 entry tracks of IFT-A (CHE-11), average docking location 71 ± 26 nm, average measured pause time is 1.2 ± 0.2 s. **(D)** Distribution of binned average velocities (solid line) along the length of the cilium for BBSome (yellow), IFT-B (blue) and IFT-A (red). **(E)** The shape of the distribution, binned along the ciliary length, obtained from moving single-molecule localizations of entry events corresponding to BBSome (yellow), IFT-B (blue) and IFT-A (red). Shaded areas (in D and E) indicate the errors. Average values and errors are estimated using bootstrapping.

## Discussion

In most ciliated systems, cilia protrude directly from the cell soma which makes it challenging to visualize the dynamics of ciliary proteins in the cell soma and near the ciliary base. In contrast, in the chemosensory neurons of *C. elegans*, the soma is separated from the cilium by a long dendrite, often tens of micrometers long. We utilized this particular geometry in PHA/PHB neurons, using SWIM^23^, to selectively illuminate (and photobleach) a small section of the neuron and observe how different IFT components move across the illuminated section. Using this approach, we visualized how IFT components traverse from the soma to the ciliary base and how they assemble into anterograde IFT trains at the ciliary base to enter the cilium.

### Diffusion plays a key role in transporting intact BBSome, IFT-B and IFT-A subcomplexes across the dendrite

First, we investigated whether IFT proteins reach the ciliary base by random diffusion or if they are specifically transported there by active transport. We found that all investigated IFT components undergo diffusion in the dendrite, with BBS-2 (BBSome) and IFT motors moving exclusively via diffusion, whereas OSM-6 (IFT-B) and CHE-11 (IFT-A) also being transported in a directed manner, as part of vesicles (discussed later). The diffusion coefficients we extracted for BBSome, IFT-A and IFT-B subunits diffusing in the dendrites are consistent with these subunits being part of the respective (BBSome, IFT-A, and IFT-B) subcomplexes and being transported as intact subcomplexes (Figure 7A). This observation is in line with several earlier findings. It has been shown before that (i) functionally related IFT proteins show similar FRAP recovery rates at the ciliary base indicating shared recruitment dynamics^8,9^, (ii) some IFT proteins cannot be expressed or purified without their subcomplex partners^36,37^ and (iii) IFT proteins mislocalize when subcomplex partners are mutated ^8,38^. It is worth to note that the IFT-B complex is composed of two subcomplexes, IFTB1 and IFTB2, which showed slightly different recruitment dynamics in a FRAP study^8^. While in this study we assume OSM-6 represents dynamics of complete IFT-B subcomplexes, it needs to be tested whether IFT-B1 and IFT-B2 always form a complex in dendrite and cilium.

**Figure 7:**
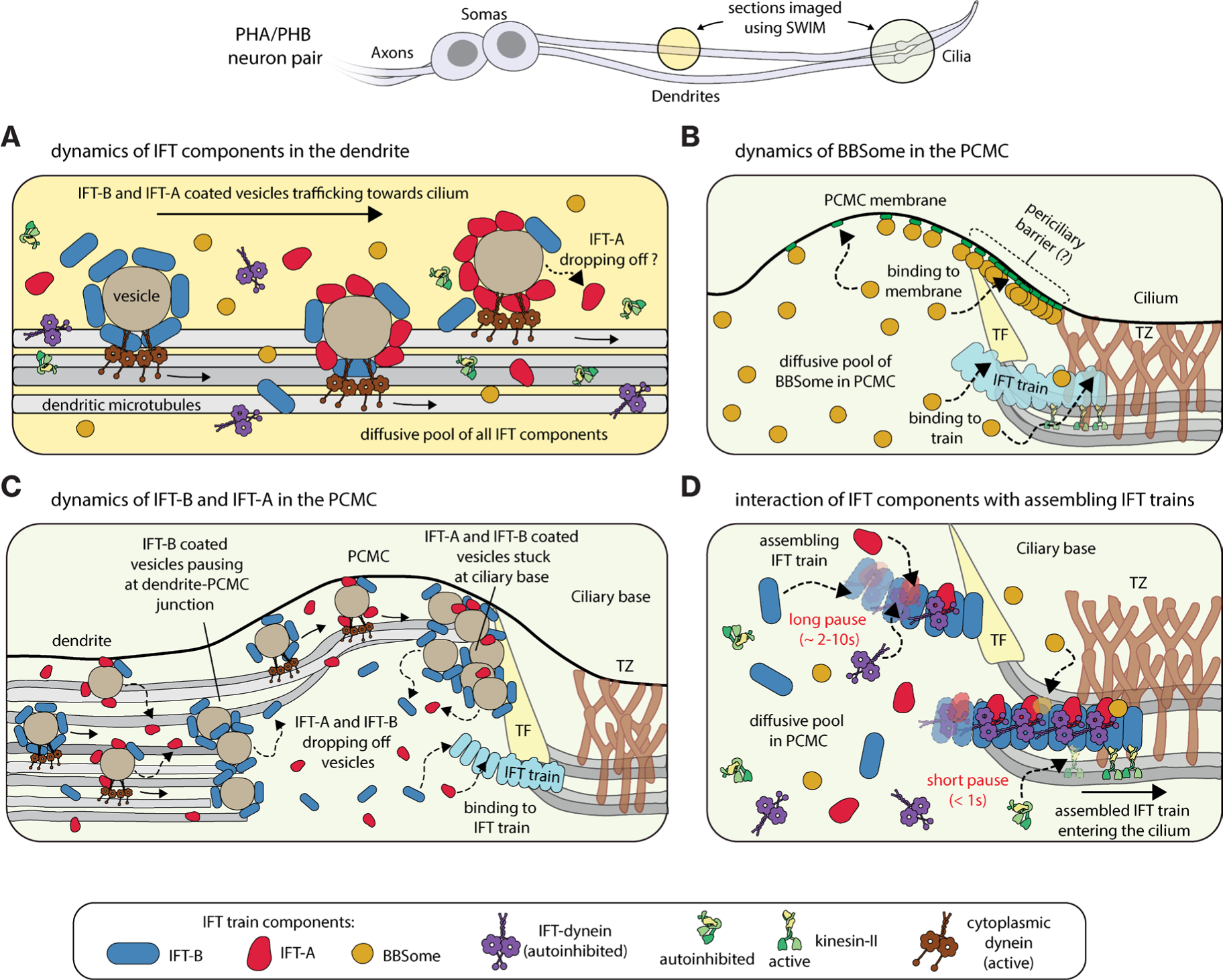
Illustration of the dynamics of IFT components as they traffic across the dendrite, organize at the PCMC and enter the cilium. **(A)** BBSomes, IFT dynein and kinesin-II move across the dendrite purely via diffusion, while IFT-B and IFT-A are transported in a directed manner via vesicles. **(B)** The diffusive pool of BBSomes at the PCMC either associates with anterograde IFT trains parked at the ciliary base or with the PCMC membrane, primarily in the periciliary barrier (region between the TZ and the TF). **(C)** The dynamics of IFT-A and IFT-B-coated vesicles at the PCMC. IFT-B-coated vesicles stall temporarily at the dendrite-PCMC junction before parking for a much longer duration at the ciliary base. IFT-A coated vesicles also park at the ciliary base, but do not stall at the dendrite-PCMC junction. Individual IFT subcomplexes fall off the vesicles as they move towards the ciliary base. IFT-A dissociates from vesicles more frequently than IFT-B resulting in a much larger pool of vesicle-associated IFT-B than IFT-A at the ciliary base. **(D)** IFT components load onto assembling anterograde IFT trains from a diffusive pool in the PCMC. BBSome and kinesin-II associate with IFT trains only during the late stages of assembly while IFT-A, IFT-B and IFT-dynein associate with IFT trains throughout the period of assembly.

### At the ciliary base, BBSomes associate with the PCMC membrane or with assembling IFT trains

Visualization of the single-particle dynamics of BBSomes diffusing near the ciliary base revealed that they either associate with the PCMC membrane, or bind to assembling anterograde IFT trains that, after assembly is completed, transport the BBSomes into the cilium (Figure 3A-3D; Figure 7B). Membrane-associations occur throughout the PCMC but are enriched proximal to the TZ in an ∼300 nm wide region (Figure 3D). It has been shown before that BBSomes are recruited to the membrane by ARL-6/BBS-3, switching from a closed autoinhibited state in solution to an open membrane-bound conformation^29,30^. ARL-6/BBS-3 was found to be enriched at the ciliary base, close to the transitional fibers in mammalian cells^32,33^ and *Chlamydomonas* ^39^. Furthermore, recent studies have revealed that BBSomes and ARL-6/BBS-3 associate with retrograde IFT trains to remove activated GPCRs from cilia^40^. The activated GPCRs are driven across the TZ into an intermediate compartment located between the TZ and a nearly impassable periciliary barrier, where they reside before being recycled into the cilium^35,40^. Based on our observations and these previous findings we hypothesize that, in *C. elegans*, BBSomes are recruited to the PCMC membrane by ARL-6/BBS-3, which is likely enriched at the proposed intermediate compartment. We also observed that retrograde-moving BBSomes cross the TZ and localize in the membrane-bound pool (Figure 3C), suggesting that BBSomes plays a key role in ciliary gating of membrane proteins at the ciliary base. To our surprise, BBSomes entering the cilium pause only for a short duration and the localization profile of the entering BBSome is distinct from the membrane-bound pool (Figure 3D-3F). This suggests that BBSomes associating with IFT trains directly originate from the cytosolic pool and not from the ARL-6/BBS-3-associated membrane-bound pool. In *Chlamydomonas* it has been shown that ARL-6/BBS-3 is required for recruiting BBSomes to the basal body but not for ciliary entry^41^. This suggest that BBSomes are present in two separate pools at the ciliary base: (i) a membrane-bound pool primarily localized between the TZ and the transition fibers, which likely forms the periciliary barrier^35,40^, and (ii) a pool associating with stationary anterograde trains assembling at the ciliary base. Further single-particle studies of the BBSome in conjunction with ARL-6/BBS-3 and transition fiber proteins, like FBF-1, could reveal a more detailed picture on the functional role of BBSomes at the ciliary base.

### IFT-B and IFT-A are carried via vesicles to the PCMC, where they dissociate and form a diffusive pool

In contrast to BBSomes and IFT motors, IFT-A and IFT-B are transported in an active manner from the soma to the PCMC, most likely, in the form of vesicles coated with these IFT subcomplexes (Figures 2D-2I; Figure 7A), with previous studies also showing that several IFT-B and IFT-A proteins cluster around periciliary vesicles^16–19^. Furthermore, in *C. elegans*, several ciliary transmembrane proteins like OCR-2^23,26^ and OSTA-1^42^ as well as cytosolic proteins like guanylyl cyclase GCY-12^43^, tubulin glutamylase TTLL-11^44^ have been shown to undergo directed vesicular trafficking across the dendrite and these periciliary vesicles appear to accumulate in the PCMC^45,46^. Thus, diverse cellular components are targeted in a specific manner to dendritic ends, transported at different rates (whenever characterized^23,43^), with IFT proteins likely acting as vesicular coat proteins. Based on this, we hypothesize that IFT-B and IFT-A proteins play a key extraciliary role in selective sorting and transport of proteins to the PCMC.

We observed that the dynamics of IFT-A and IFT-B coated vesicles entering the PCMC are not completely the same. IFT-B-coated vesicles often paused briefly at the dendrite-PCMC junction, located ∼0.9 *μm* from the ciliary base, before moving to the ciliary base where they stalled for a longer duration (Figure 7C). The PCMC membrane is known to be a separate compartment from the dendrite^47^ and is enriched with cilium-related sensory signaling proteins as well as regulators of endocytosis and membrane trafficking^45,48,49^. The mechanism of this compartmentalization is not well understood and it is unclear whether IFT-B-coated vesicles perform a functional role while stalling at the compartment junction. The final “parking” destination for most IFT-B-coated vesicles is adjacent to the TZ at the ciliary base^50^. We note that the parking location of the vesicles is on the dendritic side of the intermediate compartment, where the membrane-bound pool of BBSome localizes, suggesting that these vesicles cannot cross the periciliary barrier formed by the transition fibers^5,40^. Furthermore, this parking location of IFT-B-coated vesicles is slightly proximal to the region where individual IFT-B subcomplexes associate with stationary IFT trains. In line with this observation, in *Chlamydomonas*, IFT-B has been shown to have distinct anterior and posterior pools at the ciliary base, with IFT-A primarily colocalizing with only the anterior pool^51^. This suggests that the organization of IFT-B subcomplexes at the ciliary base into two distinct pools, corresponding to periciliary vesicles and assembling IFT trains, might be similar in *C. elegans* and *Chlamydomonas*. In contrast to IFT-B, IFT-A-coated vesicles frequently stall in the dendrite (around the region ∼1.5 *μm* from the ciliary base) as well as along the PCMC membrane and the ciliary base, but far less at the dendrite-PCMC junction. In dual-color experiments, IFT-A and IFT-B co-localized in the majority of vesicles, although vesicles containing exclusively IFT-A or IFT-B were also observed. This suggests that there is a rich compositional diversity of periciliary vesicles and it is plausible the IFT-A and IFT-B proteins are implicated with selective sorting of vesicles at different locations in the PCMC. We also observed that both IFT-B and IFT-A-coated vesicles move along the periphery of the PCMC, indicating that the dendritic microtubules extending into the PCMC are organized along the PCMC membrane and likely facilitate the interaction of periciliary vesicles with the membrane.

We observed that individual IFT-A and IFT-B subcomplexes detach from IFT-coated vesicles moving into the PCMC from the dendrite, forming a pool of IFT subcomplexes diffusing in the PCMC (Figure 7C), as well as the dendrite (Figure 7A). IFT-B subcomplexes mostly dissociate from vesicles after they stall at the ciliary base while IFT-A subcomplexes rapidly detach from vesicles while they are moving towards the ciliary base. This results in the significant enrichment of IFT-B at the ciliary base in comparison to IFT-A, as observed in the ensemble fluorescence-intensity profiles (relative to the pool in the TZ; Figure 1G). IFT-B and IFT-A entry events primarily originate from the diffusive pool of IFT subcomplexes. We never observed IFT-B entry events directly originating from the large number of vesicles stalled at the ciliary base, although on rare occasions, IFT-A originating from vesicles does appear to enter the cilium (Figure 4F). A recent structural study has shown that the IFT-A domains required to coat periciliary vesicles are shielded when IFT-A polymerizes on IFT trains^52^, suggesting that IFT-A cannot bind with vesicles and IFT trains at the same time. Insights on IFT-B in this regard are still lacking. Overall, our findings suggest IFT-train subcomplexes enter the cilium mostly from a diffusive PCMC pool, replenished with individual IFT-train subcomplexes that dissociate from continuously arriving IFT-B and IFT-A coated vesicles. Thus, also in *C.* elegans cilia, where active transport does play a role in transporting the IFT subcomplexes along the dendrite towards the ciliary base, association of the subcomplexes with assembling IFT trains primarily occurs via a diffusion-to-capture mechanism, as proposed before in *Chlamydomonas*^20^.

### IFT components associate with assembling IFT trains in subsequent stages

We observed that individual BBSome, IFT-B and IFT-A subcomplexes enter the cilium after first pausing for a while, apparently being incorporated into immobile, assembling IFT trains at the ciliary base (Figure 7D), similar to what we observed before for IFT-dynein^11^. A recent electron-microscopy study in *Chlamydomonas* showed that anterograde IFT trains in different stages of assembly lined-up at the ciliary base, tethered at the distal end of the TZ, with IFT-A and IFT-dynein linearly oligomerizing from front to back on an IFT-B scaffold^10^. This is in line with our finding that on average IFT-B pauses for the longest (>9 s), followed by IFT-A (∼2-11 s) and IFT-dynein (∼2.7-8 s^11^), as they bind to immobile assembling trains. In contrast, BBSomes only pause briefly (∼0.4-0.7 s), similar to what we have observed before for kinesin-II^11^, indicating that BBSomes associate to anterograde IFT trains that are completely assembled or at a relatively late stage of assembly. This suggests that assembling and fully-assembled IFT trains have different affinities for several IFT components. Indeed, structural differences between assembling and moving trains have been reported^10^ and one hypothesis suggests involvement of an unknown exogenous factor that assists train polymerization, while also locking the train in an arrested conformation^53^. Finally, single-molecule localization maps in the proximal part of the cilium reveal that IFT-B is located closer to the axoneme than IFT-A and BBSomes, which are directed towards and might associate with the ciliary membrane. This observation complements structural studies of anterograde IFT trains^7,53,54^ and highlights the role of IFT-A in mediating entry of signaling receptors into cilia^55–57^. While the BBSome does not appear to play a role in ciliary entry of membrane proteins, it has been suggested to be involved in anterograde IFT of membrane proteins^58^, which would explain its close proximity to the ciliary membrane.

In summary, with our single-molecule imaging approach, we have provided a dynamic view on how different ciliary components reach and organize at the PCMC, and associate with anterograde IFT trains to enter the cilium in *C. elegans*.

## Methods

### C. elegans strains

The worm strains used in this study are listed in Table S1. The strains used in this study have been generated using Mos-1 mediated single-copy insertion^59^ and CRISPR/Cas9 genome editing^60^. Maintenance was performed using standard *C. elegans* techniques^61^, on NGM plates, seeded with HB101 *E. coli*.

### Fluorescence microscopy

Images were acquired using a custom-built laser-illuminated widefield fluorescence microscope, as described previously^62^. Briefly, optical imaging was performed using an inverted microscope body (Nikon Ti E) with a 100x oil immersion objective (Nikon, CFI Apo TIRF 100x, N.A.: 1.49) in combination with an EMCDD camera (Andor, iXon 897) controlled using MicroManager software (v1.4). 491 nm and 561 nm DPSS lasers (Cobolt Calypso and Cobolt Jive, 50 mW) were used for laser illumination. Laser power was adjusted using an acousto-optic tuneable filter (AOTF, AA Optoelectronics). For performing small-window illumination microscopy (SWIM)^23^, the beam diameter was changed using an iris diaphragm (Thorlabs, SM1D12, ø 0.8-12 mm) mounted between the rotating diffuser and the epi lens, at a distance equal to the focal length of the latter. The full beam width in the sample was ∼30 μm (2*σ* of the Gaussian width). The aperture size of the diaphragm was adjusted manually to change the width of the beam, with a minimum beam width of ∼7 μm at the sample, when the diaphragm is closed to a minimum diameter of 0.8 mm. Fluorescence light was separated from the excitation light using a dichroic mirror (ZT 405/488/561; Chroma) and emission filters (525/50 and 630/92 for collecting fluorescence excited by 491 nm and 561 nm, respectively; Chroma).

For imaging live *C. elegans*, young adult hermaphrodite worms, having PHA/PHB dendrites ∼50 μm long, were sedated in 5 mM levamisole in M9, sandwiched between an agarose pad (2% agarose in M9) and a coverslip and mounted on a microscope^62^. To perform SWIM, a small excitation window (width ranging between 7-15 μm) was used to illuminate fluorescent molecules (IFT proteins labelled with eGFP, mCherry or wrmScarlet) in a small region of PHA/PHB neurons, to image protein dynamics in either the cilia or the dendrites (see Supplementary Videos).

Samples were typically imaged at 5.3x preamplifier gain and 300 EM gain with 10MHz ADC readout and pixel size of acquired images was 80 nm × 80 nm. To image ensemble dynamics and obtain average fluorescence-intensity profiles, a cilia pair was imaged with a low intensity 491 nm or 561 nm beam (∼0.1 mW/mm^2^ in the centre of the beam) for 100 frames at 6.6 fps. To perform single-molecule imaging, sample was illuminated with a high intensity 491 nm or 561 nm beam (∼10 mW/mm^2^ in the centre of the beam). High-intensity illumination bleaches almost the whole pool of fluorescing molecules in the illuminated region of the sample, allowing visualization of “fresh”, not-yet-bleached single molecules entering the small illuminated region. Sample was imaged for 10-45 mins and image acquisition rate ranged between 4-100 fps, depending on the motility feature being visualized (diffusion, static or directed motion) and the fluorescence protein being imaged (eGFP, mCherry or wrmScarlet). For imaging diffusion of IFT components, the framerates were: BBSome (eGFP::BBS-2) 50 fps; IFT-B (OSM-6::eGFP) 50-100 fps; IFT-A (CHE-11::mCherry) ∼30 fps; IFT dynein (XBX-1::eGFP) 50-100 fps; kinesin-II (KAP-1::eGFP) 50-100 fps; OCR-2 (OCR-2::eGFP) 20-50 fps. Directed transport of packets of IFT-B and IFT-A in the dendrites was imaged at 4-20 fps (typically at 6.6 fps). Dynamics of single-particles at ciliary base was imaged at 6.6 fps for IFT-A and at 6.6 fps or 20 fps for BBSome and IFT-B.

For alternating dual-colour imaging of eGFP (OSM-6::eGFP; IFT-B) and wrmScarlet (CHE-11::wrmScarlet; IFT-A) an Arduino-compatible board (Adafruit ItsyBitsy M4) was utilized to enable communication between the camera and the AOTF module such that we alternate between 491 nm and 561 nm laser lines every 75ms, as described in Zhang et al^27^. The emitted light was separated by a two-way beam splitter, collecting signal emanating from eGFP and wrmScarlet on different regions of the same camera chip.

### Image analysis

#### Intensity profiles

First, we generated a time-averaged projection of 100 frames of the cilium pair imaged at low laser intensity (see example projections in Figures 1C-1E upper panel and corresponding movies in Video S1)). An intensity profile along the length of a cilium was made by measuring the intensity along a manually drawn segmented line (typical linewidth 9 pixels), using ImageJ. The background intensity was measured by placing a line of the same dimensions next to the cilium, which was subsequently subtracted from the raw signal to obtain the background-corrected value. The peak at the ciliary base was manually selected as 0 *μm*, to align the intensity profiles of cilia from several worms. Intensity was normalized to the value at peak intensity. Average cilium-intensity profiles were obtained by averaging the intensity profiles of multiple individual cilia with error provided by the S.D.

#### Single-molecule tracking considerations

Single-molecule tracking was only performed on those worms that did not show micron-range movement during the entire acquisition time (ranging between 8-45 min). Movement of the worm or image drift in the nanometer range could not be accounted for and could have a minor impact on the numbers we acquire from our analysis. The localization precision of individual fluorescing molecules is likely in the range of 40 nm (2*σ*), as estimated for surface-bound eGFP in our experimental set-up^22^, although it is slightly higher for moving molecules due to motion blur^63^.

#### Tracking and spline fitting

Single-particle events were tracked using a MATLAB-based software, FIESTA (version 1.6.0) ^64^. Tracks, corresponding to an event, contain information regarding time (*t*_*i*_), x and y coordinates (*x*_*i*_, *y*_*i*_) and distance moved (*d*_*i*_), for every frame *i*. Minimum track length was 12 frames. Erroneous tracks, primarily caused by two (or more) single-molecule events too close to discriminate, were also excluded from further analysis (though used for fitting the spline). While retrograde events were observed during imaging and reported about anecdotally for BBSome (Figure 3C), they were not tracked and analysed further in this study. The strategy employed in SWIM selects for “fresh”, not-yet-bleached particles entering the cilium and for imaging single-molecule retrograde events in a statistically robust manner we would need to use a different imaging strategy.

#### Transformation to ciliary coordinates

A ciliary coordinate system was defined by interpolating a cubic spline on a segmented line drawn along the long axis of the imaged dendrite/cilium, visualized via the single-molecule localizations obtained from tracking (see Figure S3A). Tracks excluded from further analysis were also included in visualizing the ciliary structure. A reference point was picked at the base of the characteristic “bone-shaped” structure, at the ciliary base. All single-molecule localizations were transformed from x- and y-coordinates (*x*_*i*_, *y*_*i*_) to ciliary coordinates (*c*_∥_*i*_, *c*_⊥_*i*_), with *c*_∥_ the distance from the reference point along the spline and *c*_⊥_ the distance perpendicular to the spline (*c*_⊥_), as illustrated in Figure 1C.

Since the characteristic “bone-shaped” structure at the ciliary base was slightly shifted for the different IFT components, the reference points were adjusted such that the shape of the velocity profiles along the cilium length (Figure 6D) and the shape of the localization distribution (Figure 6E) were aligned.

#### Classification of directed transport and pausing

To obtain a quantitative measure for the directedness of the motion, we used an MSD-based approach to extract the anomalous exponent (α) from *MSD*(*τ*) = 2Γ*τ*^*⍺*^(where Γ is the generalized transport coefficient and *τ* is the time lag) along the track, in the direction of motion. α is a measure of the directedness of the motion, α = 2 for purely directed motion, α = 1 for purely diffusive motion and α < 1 for sub-diffusion or pausing. For each datapoint (*c*_∥_*i*_), we calculate *⍺* in the direction parallel to the spline (*⍺*_∥_*i*_), using a windowed Mean Square Displacement classifier (wMSDc) approach, described in Danné et al. ^65^. α was calculated analytically, using the following equation:

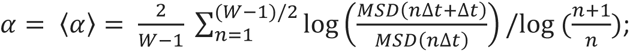

keeping a fixed window (W = 12 frames). Due to the size of the window, all tracks shorter than 12 frames were removed from the analysis. Datapoints with *⍺*_∥_*i*_ > 1.2 are classified as directed and *⍺*_∥_*i*_ < 1 are classified as static.

#### Diffusivity measurement

The covariance-based estimator (CVE) was used to estimate the diffusion coefficient of diffusive tracks along the longitudinal axis of the dendrite, using the following equation: 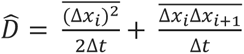; where Δ*x_i_* = *c*_∥_*i*+1_, *c*_∥_*i*_ and Δ*t* is the frame time. CVE is optimal for estimating diffusion coefficients from short, noisy trajectories of freely diffusing particles in an unbiased and regression-free manner ^24^.

#### Velocity measurement

Before calculating the point-to-point velocity, the tracks were smoothened by rolling frame averaging over 10 consecutive time frames, to reduce the contribution due to localization error (typically estimated to be between 10-40 nm, depending on the brightness of the tracked object). The point-to-point velocity at a given localization (*x*_*i*_, *y*_*i*_) was calculated using the following equation: 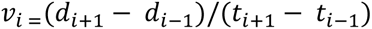.

Only moving data points (see details of classification method below) are displayed in the Figure 5F and Figure 6D, where we calculated the average velocity and error of the velocity data at bins of 100nm along the long axis of the cilia/dendrite, using bootstrapping.

#### Time-lag between dendritic packets

The time-lag between subsequent packets moving across the dendrite was extracted by manually selecting events on a kymograph. The characteristic time lag (*τ*) was obtained by least-squares fitting of the cumulative distribution function (LSF-CDF), where the time lag distribution was used to generate a cumulative probability distribution that was fitted with the CDF: *y* = 1 − *e*^−*x*⁄*τ*^. The average value and error were estimated using bootstrapping.

#### Measurement of packet intensity

The intensity of individual dendritic packets was estimated by manually drawing a box on the dendrite, with dimensions sufficient to capture all the light emitted by a single packet, positioned right at the edge of the illuminated spot, using ImageJ. Since the square was placed at the edge of the illumination spot, we minimized prior excitation and photobleaching which would decrease the measured intensity. The intensity of actively transported packets passing through the square was recorded. The background intensity was measured by placing a box of the same dimension outside the dendrite and this was subsequently subtracted from the raw signal to obtain the background corrected value.

#### Numerical simulations of 1D projection of 2D single-molecule localizations

Distribution of 2D localizations (y_*i*_, z_*i*_) along a transverse cross-section of the modelled hollow cylinder representing the axoneme (scheme illustrated in Figure S3B), was numerically simulated (N = 100000), with *r*_*ax*_ = 150 nm providing the width of the cylinder, *r*_*db*_ = 25 nm providing the thickness of the ring and localization precision estimated to be 40 nm (2*σ* for normal distribution). Coordinates of individual localizations were obtained as follows:

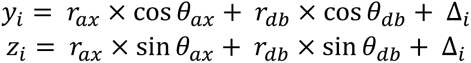

where, *θ*_*ax*_ and *θ*_*db*_ was randomly assigned to the localization and Δ_*i*_ (localization precision) was randomly picked from a normal distribution with width 40 nm (2*σ*). The distribution of *y*_*i*_, a bimodal distribution centred at 0, was fitted with a kernel density estimation which provided the 1D projection distribution of localizations along the transverse cross-section of a hollow cylinder. For simulating 2D localizations along the PCMC, we assumed a cylinder of width *r*_*PCMC*_ = 280 nm (replacing *r*_*ax*_ in the above equations), negligible thickness (*r*_*db*_ = 0 nm) and Δ_*i*_ = 100 nm.

#### Filtering events at ciliary base

Tracks at the PCMC and the cilium were classified into packet events, stuck events and entry events as follows: (i) Entry events: *c*_∥_1_ ≥ −0.3 *μm*, *c*_∥_*end*_ − *c*_∥_1_ ≥ 0.3 *μm* and max (*c*_∥_*i*_) > 0.3 *μm*; (ii) PCMC events: *c*_∥_1_ ≤ −0.5 *μm* and *c*_∥_*end*_ ≤ 0.6 *μm* and (iii) Stuck events: remaining events, where *c*_∥_1_ and *c*_∥_*end*_ are the distance along the spline for the first and last data point of a given track. For BBSome tracks, each event which was not an entry event was classified as a stuck event.

#### Measurement of docking location

Docking location of a single-molecule entry track was defined as the averaged coordinates (*c*_∥_, *c*_⊥_) of the first 2 frames.

#### Measured Pause time

Measured pause time, *t*_*p*_*m*_, of a single-molecule entry track was defined as the time taken (interpolated) to move the first 100nm. Tracks shorter than 300 nm were discarded from the distribution.

#### Bleach time fit

The characteristic bleach time *t*_*bleach*_ for a given fluorescently-labelled protein in the cilia was obtained from the exponential fit (*e*^−*x*/*tbleach*^) to the decay in the fluorescence intensity (normalized) over time, as a result of bleaching induced by the high intensity laser exposure (as shown for OSM-6::eGFP in Figure S5B). The fluorescence intensity was measured on ImageJ by manually selecting a region containing fluorescent signal and a region next to it within the illuminated area, for background correction.

#### Numerical simulations of single-molecule trajectories

To determine the impact of bleaching on the measured pause time, entry events were numerically simulated (scheme illustrated in Supplementary Figure 5A), similar to a previous study^11^. Simulated tracks were allowed to dock at a given location along a 1D cilia lattice, with the docking location randomly picked from the distribution of docking locations obtained experimentally. Each track was assigned a bleach time (*t*_*b*_) and a “real” pause time (*t*_*p*_) randomly picked from exponential distributions with characteristic time *t*_*bleach*_ (obtained from experiments) and *t*_*p*_*real*_ (free parameter), respectively. The location of the simulated event was updated every Δ*t* (with Δ*t* = 150ms, as in most experiments) and the localization precision was estimated to be 40nm (standard deviation 2*σ*). At a frame I, while *t*_*i*_ < *t*_*b*_ or *D*_*i*_ < 4μm, the location of the event was updated as follows:

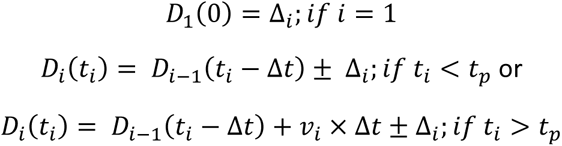

where, 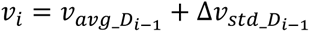 (*v*_*avg*_ is the location-dependent velocity at *D*_*i*−1_ obtained from experiments; Δ*v*_*std*_ is a randomly picked value from a normal distribution with width being the error in velocity at *D*_*i*−1_ obtained from experiments) and Δ_*i*_ is the localization precision, randomly picked from a normal distribution with width 40nm (2*σ*). For each simulation condition, with number of events N, the “real” pause time, *t*_*p*_*real*_, is varied and the measured pause time, *t*_*p*_*m*_, is recorded for tracks longer than 300nm, as described for experimental data.

#### Estimating the shape of the cilium from single-molecule localizations

Moving localizations of entry events were used to estimate the shape of the distribution of an IFT component along the cilia. The absolute distance perpendicular to the cilia spline was sampled every 1000 data points along the long axis of the cilia (shifting every 300nm for the next sample) from the single-molecule localization map of a given imaged species. The 80-percentile value obtained from the cumulative distribution function of the sampled distribution provided the width of the cilium at the location along the cilia spline where the distribution was sampled. We chose the 80-percentile value of the cumulative distribution function since this is approximately where the distribution of absolute distance perpendicular to the cilia spline peaks for sampled BBSome localizations. 70-90 percentile range of the cumulative distribution function was displayed to represent the error in the width.

#### Estimating average value and error for distributions

We used a bootstrapping method to calculate the parameters of a distribution. We randomly selected N measurements from the distribution (with replacement) and calculated the median of the resampled group. We repeated this process 1000 times, creating a bootstrapping distribution of medians. We then calculated the mean (μ) and standard deviation (σ) of this distribution and used these values to estimate the parameter and its error. In this paper, all values and errors are presented as μ ± 3σ.

#### Information on plots and figures

Kymographs were generated using the Multi Kymograph plug-in of Fiji/ImageJ. All the data was analyzed and plotted using custom written scripts on MATLAB (The Math Works, Inc., R2021a).

## Data availability

The data, and the MATLAB scripts for data analysis, visualization and numerical simulations associated with this manuscript are available on DataverseNL: https://doi.org/10.34894/A3ZUDN

## Acknowledgements

We thank current and previous members of our lab for fruitful discussions and Elizaveta Loseva for comments on the manuscript. We acknowledge financial support from the European Research Council under the European Union’s Horizon 2020 research and innovation programme (Grant agreement no. 788363; “HITSCIL”) and Marie Sklodowska-Curie Actions Postdoctoral Fellowship of the European Commission (Project no. 898006; ‘MingleIFT’, A.M.).

## Author contributions

Conceptualization and methodology, A.M., and E.J.G.P.; investigation, A.M., E.G. and W.M.; formal analysis, A.M.; resources, A.M.; visualization, A.M., writing – original draft, A.M.; writing – review & editing, A.M., and E.J.G.P.; funding acquisition, A.M., and E.J.G.P.

## Supplementary Figures

**Figure S1:**
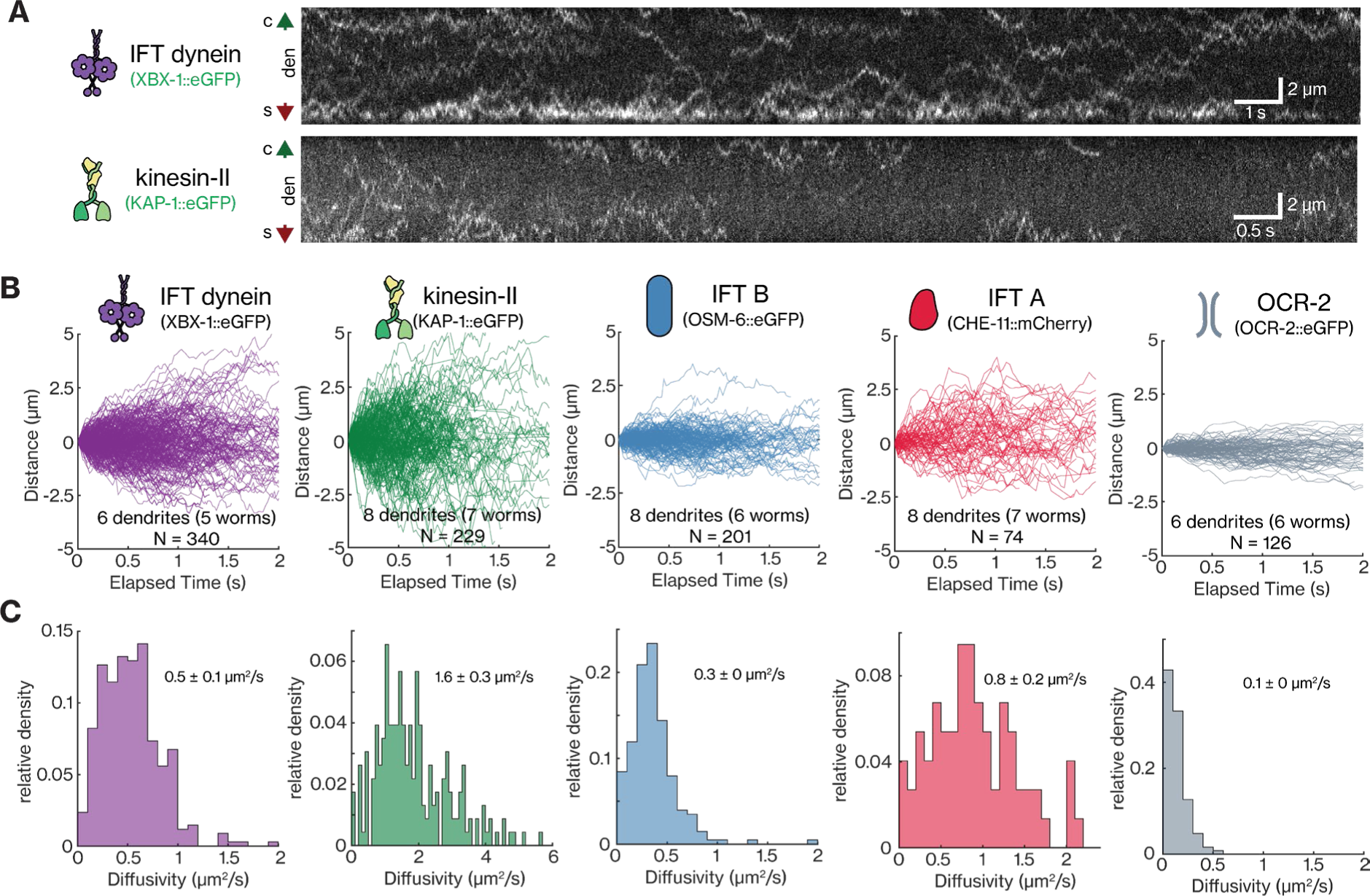
Analysis of tracked diffusive events of different IFT components diffusing in the dendrites of PHA/PHB neurons. **(A)** Representative kymograph of IFT dynein (XBX-1::eGFP; light intermediate chain subunit of IFT-dynein; top) and kinesin-II (KAP-1::eGFP; non-motor subunit of heterotrimeric kinesin-2; bottom) displays that single IFT dynein and kinesin-II motors diffuse across the dendrite (see Video S2). Green and red arrows indicate the cilium and soma direction, respectively. **(B-C)** Distance-time plots (B) and histogram of the diffusivity of individual tracks (C) for different IFT components: IFT dynein: 340 tracks in 6 dendrites, average diffusivity 0.5 ± 0.1 *μm*^2^/*s*; kinesin-II: 229 tracks in 8 dendrites, average diffusivity 1.6 ± 0.3 *μm*^2^/*s*; IFT-B (OSM-6::eGFP): 201 tracks in 8 dendrites, average diffusivity 0.3 ± 0 *μm*^2^/*s*; IFT-A (CHE-11::mCherry): 74 tracks in 8 dendrites, average diffusivity 0.8 ± 0.2 *μm*^2^/*s*; OCR-2 associated vesicles (OCR-2::eGFP): 126 tracks in 6 dendrites, average diffusivity 0.1 ± 0 *μm*^2^/*s* (data from Mitra et al.^23^). Average values and errors are estimated using bootstrapping.

**Figure S2:**
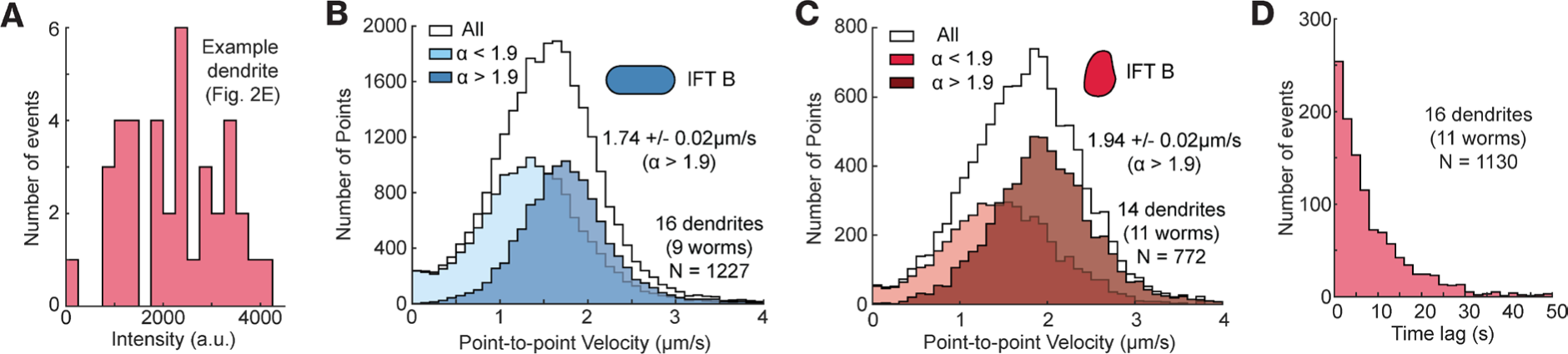
Analysis of the motion of directed packets of IFT-B and IFT-B moving from across dendrites from soma to cilia. **(A)** Histogram of intensity of directed packets of IFT-A imaged in example dendrite shown in 2D and Video S3 (right). Average values and errors are estimated using bootstrapping. **(B)** Histograms of point-to-point velocities obtained from 1277 tracks of IFT-B (OSM-6) packets from 16 dendrites. Average velocity obtained from all data points is 1.54 ± 0.01 μm/s (N = 26765; grey). For data points with α < 1.9, average velocity is 1.33 ± 0.02 μm/s (N = 15125; light blue), while, for data points with α > 1.9, it is 1.74 ± 0.02 μm/s (N = 11640; blue). **(C)** Histograms of point-to-point velocities obtained from 772 tracks of IFT-A (CHE-11) packets from 14 dendrites. Average velocity obtained from all data points is 1.76 ± 0.02 μm/s (N = 10910; grey). For data points with α < 1.9, average velocity is 1.45 ± 0.04 μm/s (N = 4738; red), while, for data points with α > 1.95, it is 1.94 ± 0.02 μm/s (N = 6172; brown). **(D)** Histogram of the time lag between subsequent IFT-A packets in 16 dendrites (N = 1130).

**Figure S3:**
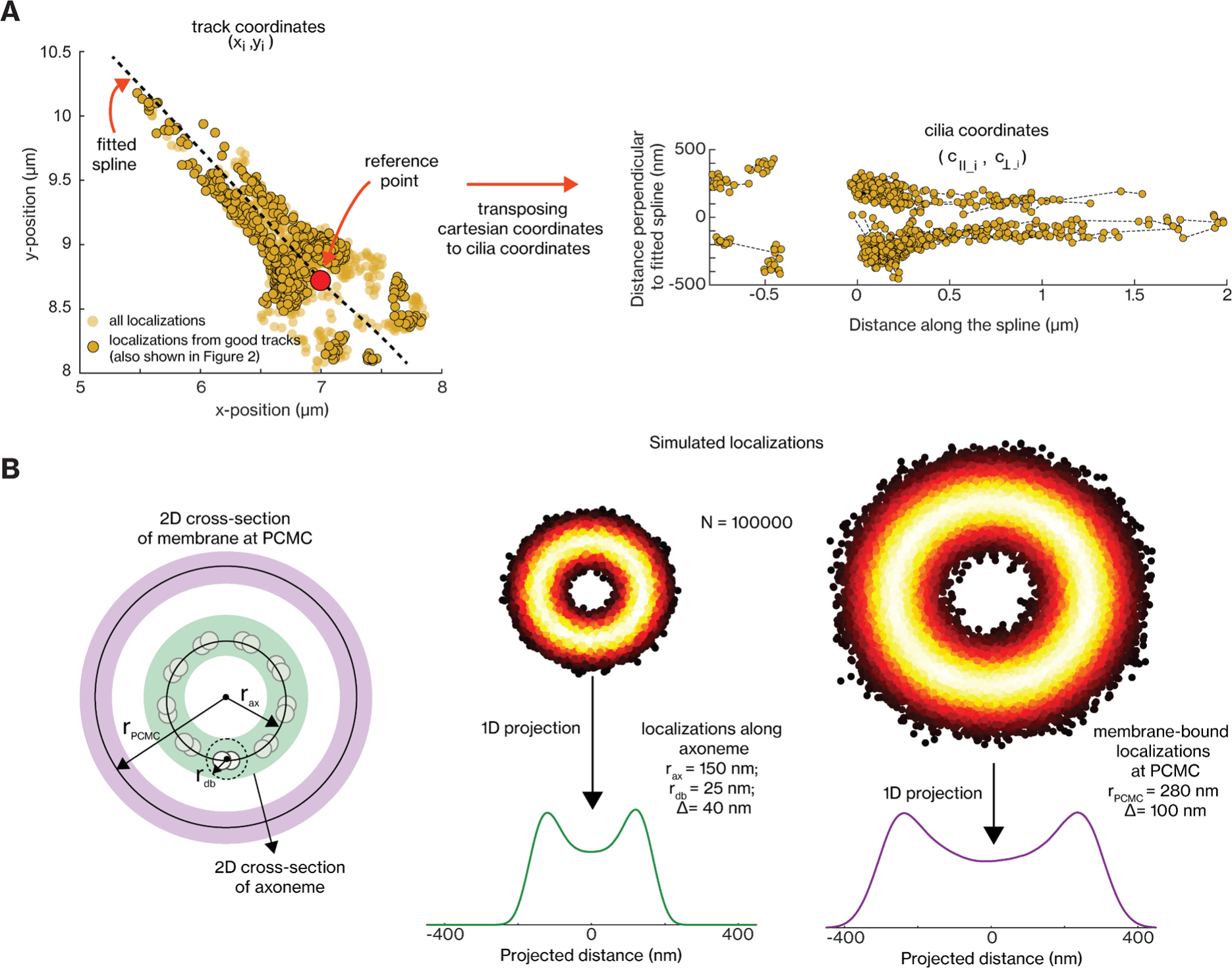
**(A)** Transposing cartesian coordinates of single-molecule tracks to general cilia coordinates. Top: x-y coordinates of all tracked kinesin-II entry events (846 single-molecule localizations from 18 tracks) in an imaged cilium. The single-molecule localizations provide the structure near the ciliary base, making it possible to draw a spline (dotted black line) roughly along the longitudinal axis of the cilium and select a reference point at the ciliary base. The cartesian coordinates of each single-molecule localization (*x*_*i*_, *y*_*i*_) can be transposed to distance perpendicular to the spline (*c*_⊥*i*_) and distance from the reference point (*c*_∥*i*_), referred to as cilia coordinates. Bottom: Single-molecule localizations replotted in cilia coordinates. **(B)** 1D projection simulations to explain the bimodal distribution (centred around 0 μm) observed for single-molecule localizations in the direction perpendicular to the fitted spline. Left: Illustration of the transverse section of the hollow axoneme (radius *r*_*ax*_; shaded in green) comprising of 9 microtubule doublets, with BBSome complexes moving along individual doublets within the radius (radius *r*_*db*_) and the transverse section occupied by membrane bound BBSome complexes at PCMC (radius *r*_*PCMC*_; shaded in purple). Right: 1D projection of the 2D cross-sectional geometry illustrated in the left. Parameters for axonemal cross-section: N = 10000; *r*_*ax*_ = 150 nm, *r*_*db*_ = 25nm and localization error Δ= 40nm. Parameters for membrane cross-section: N = 10000; *r*_*ax*_ = 280 nm and error Δ_*PCMC*_= 100nm. Δ_*PCMC*_ accounts for localization error and variability in the radius of the PCMC between worms.

**Figure S4:**
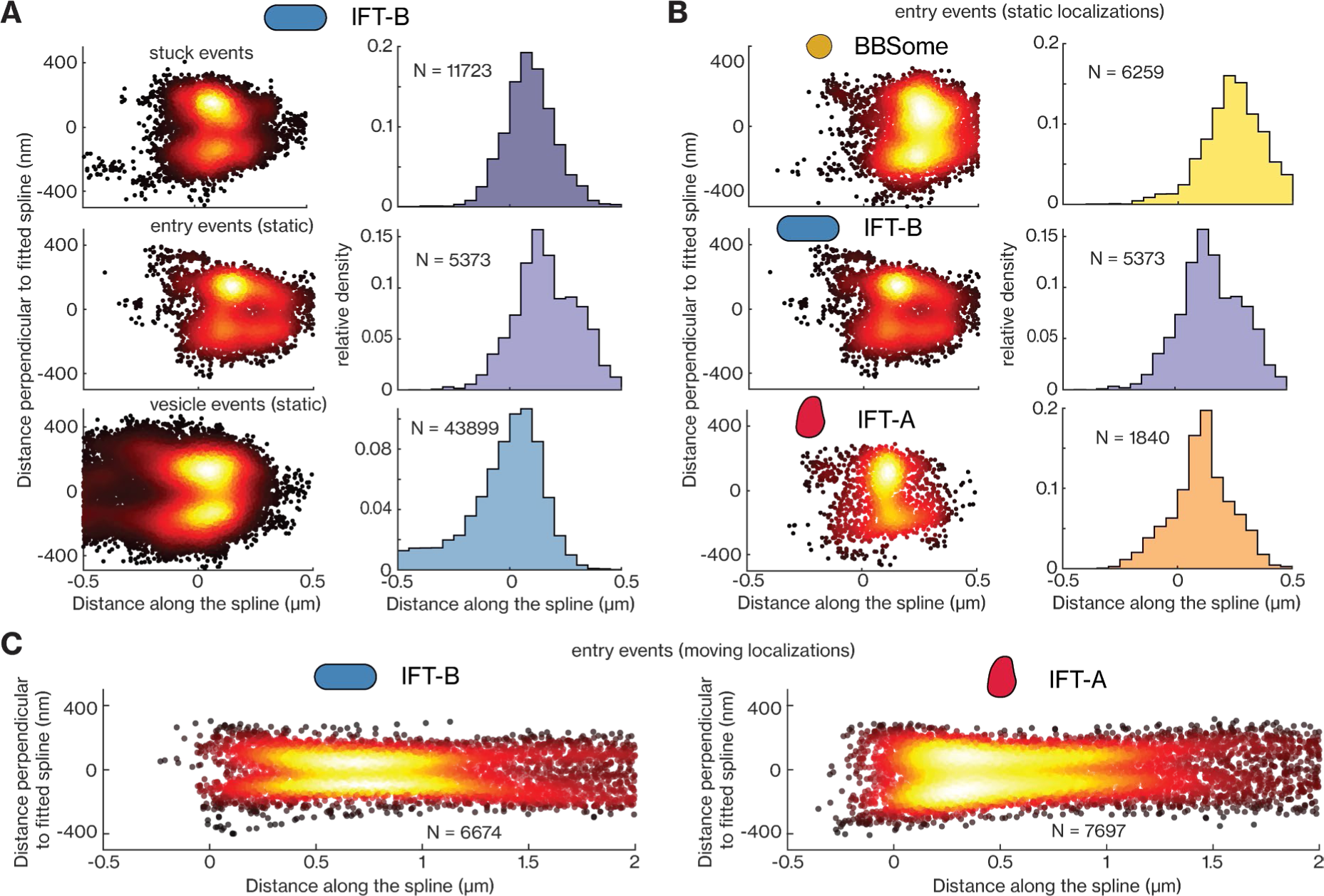
Single-particle localizations of IFT-B and IFT-A in the PCMC and ciliary base. **(A)** Super-resolution map of static single-particle localizations (left) and distribution of the localizations along the longitudinal axis of the cilia (right), obtained from tracks of IFT-B. Top: static localizations from stuck events (11723 localizations from 245 tracks); middle: static localizations from entry events (5373 localizations from 154 tracks); bottom: static localizations from vesicle events (43899 localizations from 1549 tracks). **(B)** Super-resolution map of static localizations (left) and distribution of the localizations along the longitudinal axis of the cilia (right) obtained from entry events of BBSome (top; 6259 localizations), IFT-B (middle; 5373 localizations) and IFT-A (bottom; 1840 localizations). **(C)** Super-resolution map of moving localizations of entry events for IFT-B (left; 6674 localizations) and IFT-A (right; 7697 localizations).

**Figure S5:**
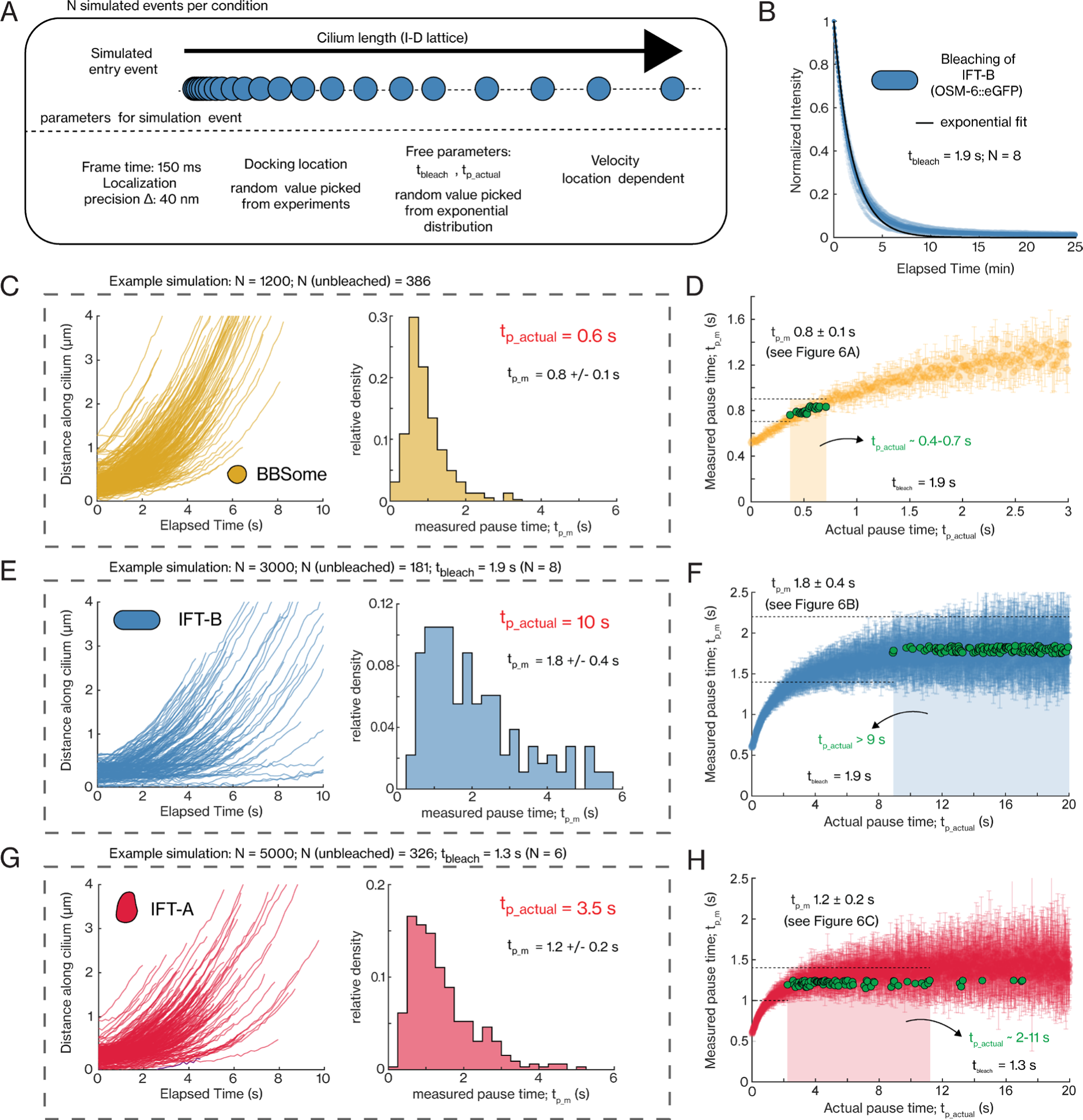
Numerical simulations to estimate actual pause time of single-molecule tracks entering cilia. **(A)** Scheme of the numerical simulation, as also performed previously^11^. Each simulated molecule is designated a bleach time (*t*_*b*_) and “actual” pause time (*t*_*p*_), randomly picked from exponential distributions with rate parameters *t*_*bleach*_ (estimated from experiments) and *t*_*p*_*actual*_ (free parameter), respectively. The molecule docks along a 1D cilium lattice, with the docking location randomly selected from experimentally measure docking locations. After every time interval, the molecule either stays in the same location (elapsed time *t* < *t*_*b*_ & *t*_*p*_), moves forward (*t* < *t*_*b*_ & *t* > *t*_*p*_) with a location dependent velocity (obtained from experiments) or beaches (*t* ≥ *t*_*b*_; end of event). N is the number of events for a given condition, frame time is 150 ms and the localization precision is 40 nm (2*σ*). **(B)** Exponential decay of the IFT-B (OSM-6::eGFP) intensity over time, upon exposure to high intensity 491 nm laser (number of cilia N = 8). The exponential fit (black line) provides a characteristic *t*_*bleach*_ = 1.9 s. **(C)** Distance-time plots of simulated BBSome entry events (N = 1200, N[unbleached] = 386) and histogram of measured pause time (average pause time *t*_*p*_*m*_ = 0.8 ± 0.1 s). **(D)** Distribution of the measured pause times, *t*_*p*_*m*_, with respect to actual pause times, *t*_*p*_*actual*_, obtained from numerical simulations of BBSome entering cilia, assuming a characteristic bleach time (*t*_*bleach*_) of 1.9 s. Each point represents the average pause time (*t*_*p*_*m*_) for a given simulated experiment. *t*_*p*_*actual*_ is in the range 0.4-0.7s for *t*_*p*_*m*_ 0.8 ± 0.1nm (experimentally obtained; Figure 6A) **(E)** Distance-time plots of simulated IFT-B entry events (N = 3000, N[unbleached] = 181) and histogram of measured pause time (average pause time *t*_*p*_*m*_ = 1.8 ± 0.4 s), for *t*_*p*_*actual*_ = 10s. **(F)** Distribution of the measured pause time, *t*_*p*_*m*_, with respect to actual pause time, *t*_*p*_*actual*_, for IFT-B, obtained from numerical simulations (using *t*_*bleach*_ = 1.9 s). *t*_*p*_*actual*_ is estimated to be >9 s for *t*_*p*_*m*_ = 1.8 ± 0.4s (experimentally obtained; Figure 6B). **(G)** Distance-time plots of simulated IFT-A entry events (N = 5000, N[unbleached] = 326) and histogram of measured pause time (average pause time *t*_*p*_*m*_ = 1.2 ± 0.2 s), for *t*_*p*_*actual*_ = 3.5 s. *t*_*bleach*_ is 1.3 s (N = 6). **(H)** Distribution of the measured pause time, *t*_*p*_*m*_, with respect to actual pause time, *t*_*p*_*actual*_, for IFT-A, obtained from numerical simulations (using *t*_*bleach*_ = 1.3 s). *t*_*p*_*actual*_ is estimated to be in the range 2-11 s for *t*_*p*_*m*_ = 1.2 ± 0.2s (experimentally obtained; Figure 6C). Average value and error are estimated using bootstrapping.

## Supplementary Tables

**Table S1:** *C. elegans* strains used in this study. Short notation is used throughout the main text and figures to increase readability.

| Strain | Genotype | Source | Short notation |
| --- | --- | --- | --- |
| PHX6589 | *bbs-2(syb6589[eGFP::bbs-2]) IV* | This study<br>(made by SunyBiotech) | BBSome (BBS-2) |
| EJP76 | *vuaSi15 [pBP36; Posm-6::osm-6::eGFP; cb-unc-119(+)] I; unc-119(ed3) III; osm-6(p811) V* | Prevo et al., 2015 | IFT-B (OSM-6) |
| EJP81 | *vuaSi24 [pBP43; Pche-11::che-11::mCherry; cb-unc-119(+)] II; unc-119(ed3) III; che-11(tm3433) V* | Prevo et al., 2015 | IFT-A (CHE-11) |
| EJP13 | *kap-1(ok676) III; vuaSi1 [pBP20; Pkap-1::kap-1::eGFP; cb-unc-119(+)] IV* | Prevo et al., 2015 | kinesin-II |
| EJP212 | *vuaSi26 [Pxbx-1::xbx-1::EGFP; cb-unc-119(+)] I; vuaSi2 [Posm-3::osm3::mCherry; cb-unc-119(+)] II; osm3(p802) IV; xbx-1(ok279) V* | Mijalkovic et al., 2017 | IFT-dynein |
| PHX8141 | *osm-6(syb7957[osm-6::eGFP]V,che-11(syb8141[che-11::wormScarlet])V* | This study<br>(made by SunyBiotech) | IFT-B and IFT-A |

**Table S2:**
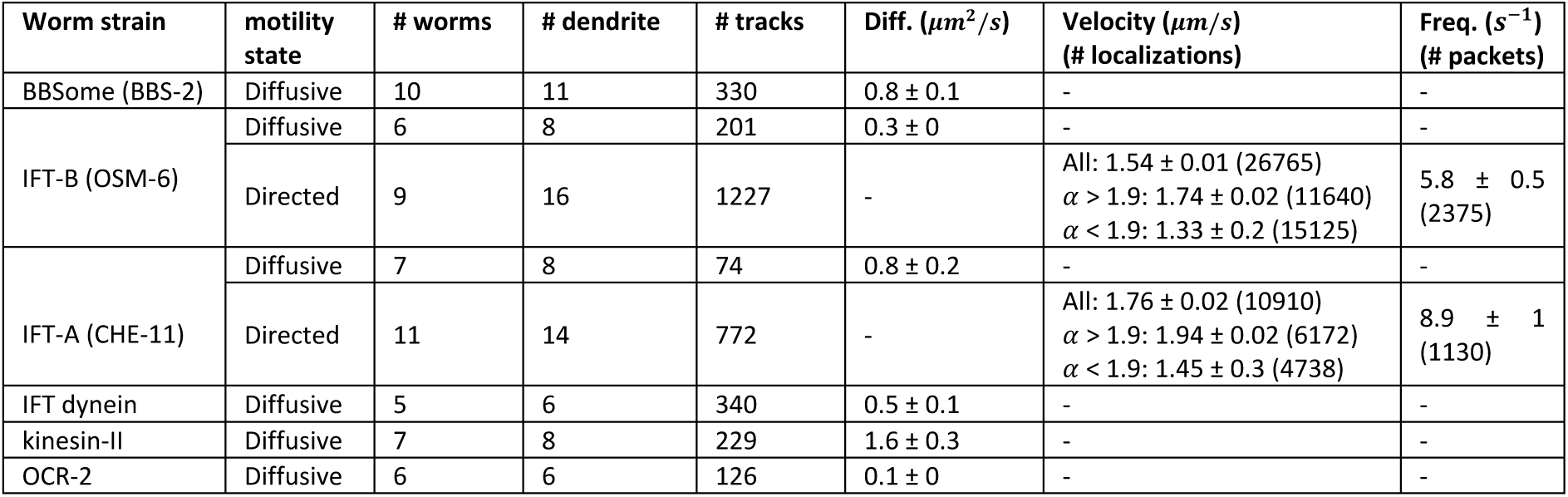
Data collected from tracking single-particle events of different IFT components in the dendrites of PHA/PHB neurons.

| Worm strain | motility state | # worms | # dendrite | # tracks | Diff. ($\mu\text{m}^2/\text{s}$) | Velocity ($\mu\text{m}/\text{s}$)<br>(# localizations) | Freq. ($\text{s}^{-1}$)<br>(# packets) |
| --- | --- | --- | --- | --- | --- | --- | --- |
| BBSome (BBS-2) | Diffusive | 10 | 11 | 330 | $0.8 \pm 0.1$ | - | - |
| IFT-B (OSM-6) | Diffusive | 6 | 8 | 201 | $0.3 \pm 0$ | - | - |
| | Directed | 9 | 16 | 1227 | - | All: $1.54 \pm 0.01$ (26765)<br>$\alpha > 1.9$: $1.74 \pm 0.02$ (11640)<br>$\alpha < 1.9$: $1.33 \pm 0.2$ (15125) | $5.8 \pm 0.5$<br>(2375) |
| IFT-A (CHE-11) | Diffusive | 7 | 8 | 74 | $0.8 \pm 0.2$ | - | - |
| | Directed | 11 | 14 | 772 | - | All: $1.76 \pm 0.02$ (10910)<br>$\alpha > 1.9$: $1.94 \pm 0.02$ (6172)<br>$\alpha < 1.9$: $1.45 \pm 0.3$ (4738) | $8.9 \pm 1$<br>(1130) |
| IFT dynein | Diffusive | 5 | 6 | 340 | $0.5 \pm 0.1$ | - | - |
| kinesin-II | Diffusive | 7 | 8 | 229 | $1.6 \pm 0.3$ | - | - |
| OCR-2 | Diffusive | 6 | 6 | 126 | $0.1 \pm 0$ | - | - |

**Table S3:** Data collected from tracking single molecule events of IFT train complexes in the PCMC and cilia of PHA/PHB neurons.

| Worm strain | # worms | # cilia | # tracks | Moving loc.<br>($\alpha > 1.2$) | Static loc.<br>($\alpha < 1$) | Docking Loc. (nm) | $t_{p,m}$ (s) | $t_{p,actual}$ (s) |
| --- | --- | --- | --- | --- | --- | --- | --- | --- |
| BBS-2<br>(BBSome) | 27 | 46 | Stuck: 456 | 437 | 13623 | - | - | - |
| | | | Entry: 367 | 11358 | 6517 | $185 \pm 23$ | $0.8 \pm 0.1$ | 0.4-0.7 |
| OSM-6<br>(IFT-B) | 16 | 30 | PCMC:1549 | 42192 | 43899 | - | - | - |
| | | | Entry: 154 | 6674 | 5373 | $98 \pm 28$ | $1.8 \pm 0.4$ | $> 9$ |
|  |  |  | Stuck: 245 | 136 | 11723 | - | - | - |
| CHE-11<br>(IFT-A) | 20 | 37 | PCMC: 727 | 11011 | 7729 | - | - | - |
| | | | Entry: 297 | 7697 | 1840 | $71 \pm 26$ | $1.2 \pm 0.2$ | 2-11 |
|  |  |  | Stuck: 264 | 409 | 6443 | - | - | - |

## Supplementary video legends

**Video S1:** Example movies of dynamics of BBSome (eGFP::BBS-2; left), IFT-B (OSM-6::eGFP; middle) and IFT-A (CHE-11:mCherry; right) in the PHA/PHB cilia of *C. elegans*, acquired at low laser intensity. Videos play at 3x real time. Time and scale bar indicated. Related to Figure 1.

**Video S2: (A-C)** Example movies (bottom panel) of single-molecule diffusive dynamics of BBSome (eGFP::BBS-2; A), IFT dynein (XBX-1::eGFP; B) and kinesin-II (KAP-1::eGFP; C) in the dendrites of PHA/PHB neurons, imaged using SWIM. Top-panel show the maximum projection corresponding to the movies, where the location of the dendrite(s) is indicated. Videos play at real time. Time and scale bar indicated. Related to Figures 2A-2C; Figure S1.

**Video S3: (A-B)** Example movies (bottom panel) of single-particle diffusive and directed dynamics of IFT-B (A) and IFT-A (B) in the dendrites of PHA/PHB neurons, imaged using SWIM. Top-panel show the maximum projection corresponding to the movies, where the location of the dendrite(s) is indicated. Videos play at 10x real time. Time and scale bar indicated. Related to Figures 2F-2I; Figure S2.

**Video S4:** Example movie obtained from dual-colour imaging of IFT-B (OSM-6::eGFP) and IFT-A (CHE-11:wrmScarlet) in the dendrites of PHA/PHB neurons. Left: green channel, middle: red channel and right: merged. Videos play at 3x real time. Time and scale bar indicated. Related to Figures 2J-2K.

**Video S5:** Example movie (right-panel) displaying single-particle dynamics of individual BBSome complexes at the PCMC and proximal part of cilia. Left-panel shows the maximum projection corresponding to the movie, where the location of the dendrites, cilia and ciliary base are indicated. Videos play at 3x real time. Time and scale bar indicated. Related to Figure 3.

**Video S6: (A)** Example movie (right-panel) displaying single-particle dynamics of individual IFT-B complexes at the PCMC and proximal part of cilia. Left-panel shows the maximum projection corresponding to the movie, where the location of the dendrites, cilia and ciliary base are indicated. Videos play at 3x real time. **(B)** Sections of the movie in A displaying example IFT-B events. Left: Directed packets of IFT-B moving from the dendrite into the PCMC. Right: Ciliary entry event and a directed packet event releasing IFT-B complexes that diffuse. Videos play at real time. In all movies, time and scale bar indicated. Related to Figure 4A-4C.

**Video S7: (A)** Example movie (right-panel) displaying single-particle dynamics of individual IFT-A complexes at the PCMC and proximal part of cilia. Left-panel shows the maximum projection corresponding to the movie, where the location of the dendrites, cilia and ciliary base are indicated. Videos play at 3x real time. **(B)** Sections of the movie in A displaying example IFT-A events. Left: Directed packets of IFT-A moving from the dendrite into the PCMC. Right: Ciliary entry events. Videos play at real time. In all movies, time and scale bar indicated. Related to Figures 4D-4F.

